# Induced pathogenicity toward open-ocean diatoms by a newly isolated filterable bacterium *Ekhidna algicida* sp. nov.

**DOI:** 10.1101/2025.09.24.678341

**Authors:** Shiri Graff van Creveld, Sacha N. Coesel, Ellen Lavoie, Vaughn Iverson, Rhonda Morales, Megan J. Schatz, Alexandra E. Jones-Kellett, Jesse McNichol, Rebecca S. Key, Jed Fuhrman, Bryndan P Durham, E. Virginia Armbrust

**Affiliations:** School of Oceanography, University of Washington, Seattle, WA 98195, USA; Molecular Analysis Facility, University of Washington, Seattle, WA 98195, USA; Department of Earth, Atmospheric C Planetary Sciences, Massachusetts Institute of Technology, Cambridge, MA 02139, USA; Biology Department, Woods Hole Oceanographic Institution, Woods Hole, MA 02543, USA; Department of Biology, St. Francis Xavier University, Antigonish, NS, Canada; Department of Biology and Genetics Institute, University of Florida, Gainesville, FL 32611, USA; Department of Biological Sciences, University of Southern California, Los Angeles, CA 90007, USA

## Abstract

Phytoplankton are the base of marine food webs. They form intricate interactions with heterotrophic bacteria ranging from mutualistic to pathogenic that together impact oceanic carbon and nutrient cycling. Our understanding of these interactions in marine environments remains primarily limited to laboratory-based studies of model organisms. Here, we report the discovery and characterization of *Ekhidna algicida* sp. nov. strain To15, isolated from the oligotrophic Pacific Ocean (16°N, 140°W) based on its algicidal effect on the pelagic diatom *Thalassiosira oceanica*. Subsequent co-culture experiments demonstrate that *E. algicida* is lethal within days to a diverse array of diatoms, with the effect mediated by bacterial exudates that remain algicidal on their own against axenic *T. oceanica* cultures. Twenty additional algicidal *Ekhidna* strains were subsequently isolated from the Pacific Ocean. Our findings reveal *E. algicida* as a potentially widespread pathogen of diatoms, that can alter microbial community composition dynamics in pelagic ecosystems.

**Teaser:** A newly discovered Pacific Ocean bacterium can kill diatoms, revealing a hidden pathogenic role in open-ocean ecosystems.

## Introduction

In the euphotic zone of the ocean, where sufficient light supports photosynthesis, marine microbial communities primarily consist of single-celled eukaryotes and bacteria that together generate and recycle nearly half the organic carbon produced on Earth each year (*1*, *2*). Diatoms are the most diverse and abundant group of eukaryotic marine phytoplankton (*3*), and their interactions with bacteria can affect biogeochemical cycles in the ocean by shaping community composition, particle aggregation, and sinking (*4*). Current understanding of mechanisms underlying these interactions is primarily derived from laboratory studies conducted with either model species isolated from different environments (*5–7*) or with co-occurring species isolated primarily from coastal, nutrient-rich habitats (*8–11*). These studies reveal that beneficial bacteria can provide diatoms with essential vitamins (*9*, *12*), or hormones that promote diatom cell cycle progression (*13*). Pathogenic bacteria can secrete enzymes such as chitinases (*14*) and proteases (*15*) or other unknown compounds that cause diatom cell lysis or disrupt the diatom cell cycle (*11*, *14*).

Microbial interactions within the open ocean may differ from those of coastal environments in part due to the low concentrations of both cells and nutrients. Pelagic microbial communities are characterized by smaller cell sizes, that increase the surface-area to volume ratio leading to more efficient nutrients uptake (*16*). The small phytoplankton cell size creates proportionally smaller phycospheres, which affect how bacteria interact with such small and dispersed phytoplankton cells (*4*, *17*, *18*). Operationally, organisms and compounds that pass through 0.2 µm pore-size filters are called the ‘dissolved fraction’ or ‘filterable organisms’. About 1-10% of open ocean heterotrophic bacteria are estimated to fall within this filterable fraction (*19*), thus standard sampling protocols routinely miss these members of the ecosystems.

Here we used live diatom cultures as both a selective “media” and as a bioindicator, to isolate open ocean algicidal bacteria. We isolated 22 filterable algicidal bacteria, 21 of which are members of the genus *Ekhidna* in the phylum Bacteroidota. Pathogenicity of Bacteroidota often rely on secretion of large proteins, such as virulence factors through the type IX secretion system (T9SS) (*20*, *21*). One *Ekhidna* strain, To15, was selected for whole genome sequencing and further co-culturing experiments to determine potential modes of pathogenicity.

On the basis of genotypic and phenotypic characteristics, we propose a novel species, *Ekhidna algicida*, sp. nov., with type strain To15^T^ (=ATCC TSD-518^T^ = DSM 119715^T^).

## Results

### *Ekhidna* are filterable algicidal bacteria found across the Pacific Ocean

To enrich for filterable diatom pathogens, seawater samples collected on two research cruises in the Tropical Pacific Ocean were filtered through 0.2 µm pore-size filters, and the resulting filtrates were added to cultures of *Thalassiosira oceanica*, *T. pseudonana* (reclassified as *Cyclotella nana* (*22*)), or mixtures of other diatoms (Table S1). The diatoms and added filtrates were incubated at sea under standard diatom growth conditions (see Methods) to propagate and enrich for potential pathogens that utilized the diatom-produced organic matter for growth. After about a week, the potentially enriched cultures were passed through 0.2 µm pore-size filters to remove the diatoms, and the filtrates were added to monoalgal diatom cultures in 24 well plates (Fig. S1A). These cultures were visually examined, and clear wells, indicative of diatom mortality and potential pathogen presence (Fig. S1A, wells B3-6), were again filtered through 0.2 µm pore-size filters. The filtrates were used for further isolation by serial dilutions into axenic *T. oceanica* or *T. pseudonana* cultures. Overall, we isolated 22 algicidal bacteria from a wide range of latitudes (23°N - 4°S), longitudes (133 - 158°W), and depths (5 - 120 m) in the Pacific Ocean (Table 1, Fig. 1A, green circles). All the isolates were pathogenic to *T. oceanica* (Fig. S1B, Table 1), and no colonies appeared on MB 1.5% agar plates. The bacterial isolates grew in half-strength liquid marine broth (50% MB, but not full-strength MB) to yield visible growth after several days; 21 of the isolates were visibly yellow, and one isolate (To107) was visibly white/beige (Fig. S1C, Table 1).

**Fig. 1.**
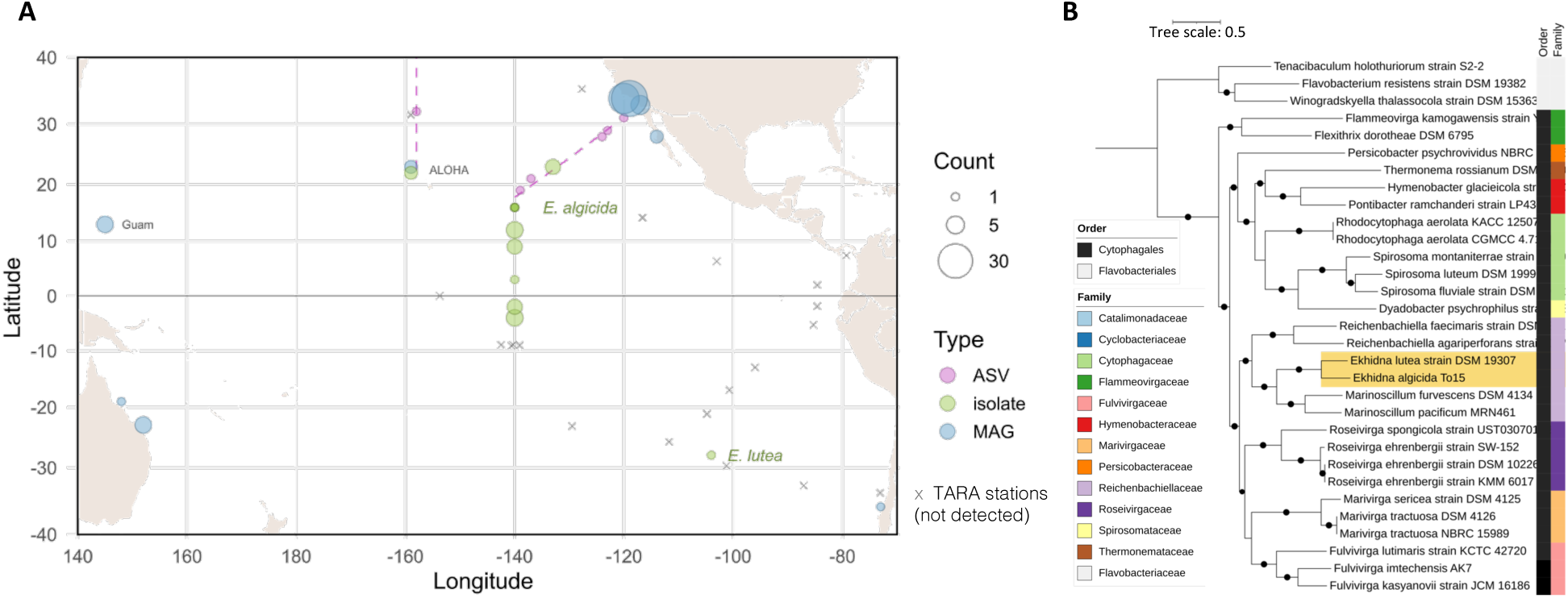
*Ekhidna* phylogeny and global distribution. **A.** Map of detection of *Ekhidna* bacteria. Colored circles indicate detection based on ASVs (purple), isolates (green) or MAGs (blue). Size of circles represents the number of detected ASVs, isolates or MAGs from a given location. Stations of the TARA expedition within the map boundary are marked with gray x’s. Gradients cruise transects (2017–2021) from which ASV data were analyzed, are marked with purple dashed lines. The *E. lutea* and *E. algicida* To15 isolates are labeled in green. **B.** A maximum likelihood phylogenetic tree made from 100 single copy genes of *Ekhidna algicida* (To15) and related bacteria. Family and order are indicated with colored vertical bars. Bootstraps of > 80 are marked with black circles, scale bar represents the number amino acid substitutions per site. *Ekhidna spp.* are highlighted with a yellow background.

**Table 1.** Algicidal *Ekhidna* isolates. Cruise identifier, latitude, longitude, depth (m), and date of sampling. The seawater filtrates were initially added to single diatoms, or mixtures of several diatoms (DM) (see Table S1). Algicidal bacteria were isolated on *T. oceanica* or *T. pseudonana*, and all are pathogenic toward *T. oceanica*. Overall, the 21 *Ekhidna* isolates exhibited 5 different 16S rDNA sequences, marked here as types A-E. Culture color when grown in 50% MB is indicated. Strain To15 is marked in bold.

| Cruise | Lat (N) | Lon (W) | Depth | Sample Date | Initial Enrichment | Isolate ID | Isolated on | *Ekhidna* 16S Type | Color | Genus |
| --- | --- | --- | --- | --- | --- | --- | --- | --- | --- | --- |
| TN412 | 23.33 | 132.7 | 5 | 2/3/23 | *T. oceanica* | To10 | *T. oceanica* | C | yellow | *Ekhidna* |
| TN412 | 23.33 | 132.7 | 5 | 2/3/23 | *T. pseudonana* | Tp2 | *T. pseudonana* | C | yellow | *Ekhidna* |
| TN412 | 23.33 | 132.7 | 5 | 2/3/23 | *T. pseudonana* | Tp3 | *T. pseudonana* | C | yellow | *Ekhidna* |
| KM24-14 | 22.49 | 158.54 | 50 | 8/23/24 | 4DM | To104 | *T. oceanica* | D | yellow | *Ekhidna* |
| KM24-14 | 22.49 | 158.54 | 50 | 8/23/24 | *T. oceanica* | To105 | *T. oceanica* | E | yellow | *Ekhidna* |
| **TN412** | **16** | **140** | **15** | **1/30/23** | **8DM** | **To15** | ***T. oceanica*** | **A** | **yellow** | ***Ekhidna*** |
| TN412 | 12.21 | 140 | 95 | 1/31/23 | 8DM | To20 | *T. oceanica* | A | yellow | *Ekhidna* |
| TN412 | 12.21 | 140 | 45 | 1/31/23 | 6DM | To23 | *T. oceanica* | A | yellow | *Ekhidna* |
| TN412 | 12.21 | 140 | 45 | 2/6/23 | 6DM | Tp4 | *T. pseudonana* | A | yellow | *Ekhidna* |
| TN412 | 12.21 | 140 | 95 | 2/6/23 | 6DM | Tp5 | *T. pseudonana* | A | yellow | *Ekhidna* |
| TN412 | 9.41 | 140 | 15 | 2/2/23 | 8DM | To12 | *T. oceanica* | A | yellow | *Ekhidna* |
| TN412 | 9.41 | 140 | 45 | 2/2/23 | 8DM | To13 | *T. oceanica* | A | yellow | *Ekhidna* |
| TN412 | 9.41 | 140 | 60 | 2/2/23 | 8DM | To14 | *T. oceanica* | B | yellow | *Ekhidna* |
| TN412 | 2.61 | 140 | 5 | 2/5/23 | *T. pseudonana* | Tp1 | *T. pseudonana* | C | yellow | *Ekhidna* |
| TN412 | -2 | 140 | 15 | 2/8/23 | 8DM | To17 | *T. oceanica* | A | yellow | *Ekhidna* |
| TN412 | -2 | 140 | 50 | 2/8/23 | 8DM | To18 | *T. oceanica* | A | yellow | *Ekhidna* |
| TN412 | -2 | 140 | 100 | 2/8/23 | 8DM | To19 | *T. oceanica* | A | yellow | *Ekhidna* |
| TN412 | -4 | 140 | 120 | 2/10/23 | *T. oceanica* | To11 | *T. oceanica* | A | yellow | *Ekhidna* |
| TN412 | -4 | 140 | 15 | 2/10/23 | *T. oceanica* | To8 | *T. oceanica* | C | yellow | *Ekhidna* |
| TN412 | -4 | 140 | 95 | 2/10/23 | *T. pseudonana* | Tp6 | *T. pseudonana* | A | yellow | *Ekhidna* |
| TN412 | -4 | 140 | 15 | 2/10/23 | *T. pseudonana* | Tp7 | *T. pseudonana* | C | yellow | *Ekhidna* |
| KM24-14 | 22.2 | 158.54 | 10 | 8/24/24 | *T. oceanica* | To107 | *T. oceanica* |  | white | *Fulvivirga* |

Full-length 16S rDNA sequences indicated that the 21 yellow isolates belong to the genus *Ekhidna,* while To107 belongs to the genus *Fulvivirga* (Table 1, Fig. S2A). Both *Ekhidna* and *Fulvivirga* are in the order Cytophagales, phylum Bacteroidetes. Within the 21 *Ekhidna* isolates, there were five unique 16S rDNA sequences, designated as sequence types A-E (Fig. S2B). Twelve isolates encode type A 16S rDNA, six encode type C, and three isolates each encode types B, D, or E (Table 1). These sequences differ from the only described *Ekhidna* species, *Ekhidna lutea* (*23*) by 4-17 nucleotides (0.4-1.6%, Fig. S2). A maximum-likelihood phylogenetic tree including the 16S rDNA of all isolated *Ekhidna* (light yellow background, Fig. S2A), and sequences derived from publicly available *Ekhidna* metagenome assembled genomes (MAGs), show the algicidal *Ekhidna* isolates, together with *E. lutea* and the other *Ekhidna* sequences in a monophyletic group (Fig. S2A). This phylogenetic organization indicates that the algicidal *Ekhidna* isolates are members of the *Ekhidna* genus, but not whether the different isolates represent different species.

We investigated the geographical distribution of *Ekhidna* bacteria by screening 4 amplicon sequence variant (ASV) datasets from previous Pacific Ocean expeditions (Gradients 1-4); *Ekhidna* 16S rDNA was detected at a total of 6 locations along the Gradients 3 (purple dashed line at 158°W) and Gradients 4 (purple diagonal dashed line) transects (Fig. 1A, purple circles). *Ekhidna* 16S rDNA was not detected in the global TARA GLOSSary dataset, as all the retrieved sequences exhibited <93% identity, and over 50 mismatches with type A *Ekhidna* 16S rRNA, and similar number of mismatches with the other *Ekhidna* isolates 16S rDNA (gray crosses in Fig. 1A mark TARA sites that fall within the map range). We searched for potential *Ekhidna* MAGs within the NCBI Genomes database and the Genome Taxonomy Database (GTDB). A total of 77 MAGs labeled as *Ekhidna* were detected (Table S2). Seventy-six of the MAGs were derived from microbiomes of diatoms, cyanobacteria, corals, sponges or kelp; one MAG (GCA_013214445.1, *Ekhidna* sp013214445) was collected from station ALOHA (22.5°N 158°W) at a depth of 4000 m, likely derived from sinking particles (Table S2). Seventy-six of the publicly available *Ekhidna* MAGs originated from the warm Pacific Ocean, between latitudes 35°N to 37°S (Fig. 1A, blue circles, Table S2); one MAG (GCA_040223195.1) was derived from the microbiome of a long-term cyanobacterium culture, isolated from a salt marsh in France.

We sequenced the whole genome of algicidal *Ekhidna* strain To15, a representative of the most abundant type A 16S rDNA, in part, to evaluate whether this strain constituted a new species. The To15 whole genome displayed 73.1% average amino-acid identity (AAI) to *E. lutea*, the closest relative within the Microbial Genomes Atlas, MiGA (*24*), suggesting that To15 is likely a previously uncharacterized species (*p*-value 0.0097). Furthermore, the average nucleotide identity (ANI) between To15 and *E. lutea* is 73.53%, well below the 95% cutoff considered to distinguish different species (*25*). A phylogenomic tree based on 100 single-copy genes (Table S3) of fully sequenced bacterial isolates (Table S4) also indicated that To15 is a novel species within the genus *Ekhidna* rather than a strain of *E. lutea* (Fig. 1B). Further analysis of To15, *E. lutea*, and *Ekhidna* MAGs revealed that To15 is most closely related to a MAG assigned to a different species from *E. lutea*, *Ekhidna* sp028821635 (GCA_006969745.1, Fig. S3), again suggesting that To15 is a previously uncharacterized species within the genus *Ekhidna*. Overall, the To15 genome, AAI and ANI analysis, and phylogenomic trees with type-material strains and MAGs indicate that the algicidal strain To15 is a previously undescribed species within the genus *Ekhidna*. According to its algicidal effect we name To15 as *Ekhidna algicida* sp. nov.

### Genomic potential of *E. algicida* To15

We analyzed the *E. algicida* To15 whole genome sequence to identify potential attributes that may underlie its observed pathogenicity. *Ekhidna algicida* is predicted to be auxotroph for nine amino acids (Lys, Phe, Ser, Thr, Pro, Val, Ile, Arg, Leu) (Table S5), similar to other parasitic and pathogenic bacteria that rely on a host for amino-acid supply and to amino-acid auxotrophies in Bacteroidota phylum (*26*). As found in other Bacteroidota, *E. algicida* encodes the five proteins of the Sec secretion system, the 15 proteins that form the T9SS (Fig. 2, Table S6) and has no indication of type 1 – 8 bacterial secretion systems. Proteins secreted by T9SS are recognized by conserved C-terminal domains (CTDs) named type A CTD and type B CTD (Fig. 2, Table S6). We detected 43 proteins with type A CTD (hmm profiles: PF18962, TIGR04183, Table S6, Fig. S4A), of which 38 also had a recognizable short N-terminal signal peptide required for the first step of secretion by the Sec system. Eight of these putatively secreted proteins were annotated as proteases: four are S8 family peptidase, three are M43 family Zn-metalloproteases and one is a C25 family cysteine peptidase (Fig. 2, Table S7). Ten proteins with type B CTDs (hmm profiles PF13585, TIGR04131) were identified (Fig. S4B, Table S7), all of which also encode for the N-terminal signal peptide; 7 of the genes encoding the type B CTDs were adjacent to genes encoding PorP proteins (Fig. S4C, Table S7), which can anchor secreted protein to the cell surface (*27*). None of the proteins with type B CTD had a functional annotation in the genome. However, HHPred analysis suggested that two of these proteins are putative grappling hook protein A (GhpA, Fig. 2, Table S7). Four additional putatively secreted proteins possess LamG-like jellyroll fold domains, a domain often found in extracellular proteins related to adhesion. Together, these results suggest that the pathogenicity of *E. algicida* may entail adhesion to target cells and secretion of proteases.

**Fig. 2.**
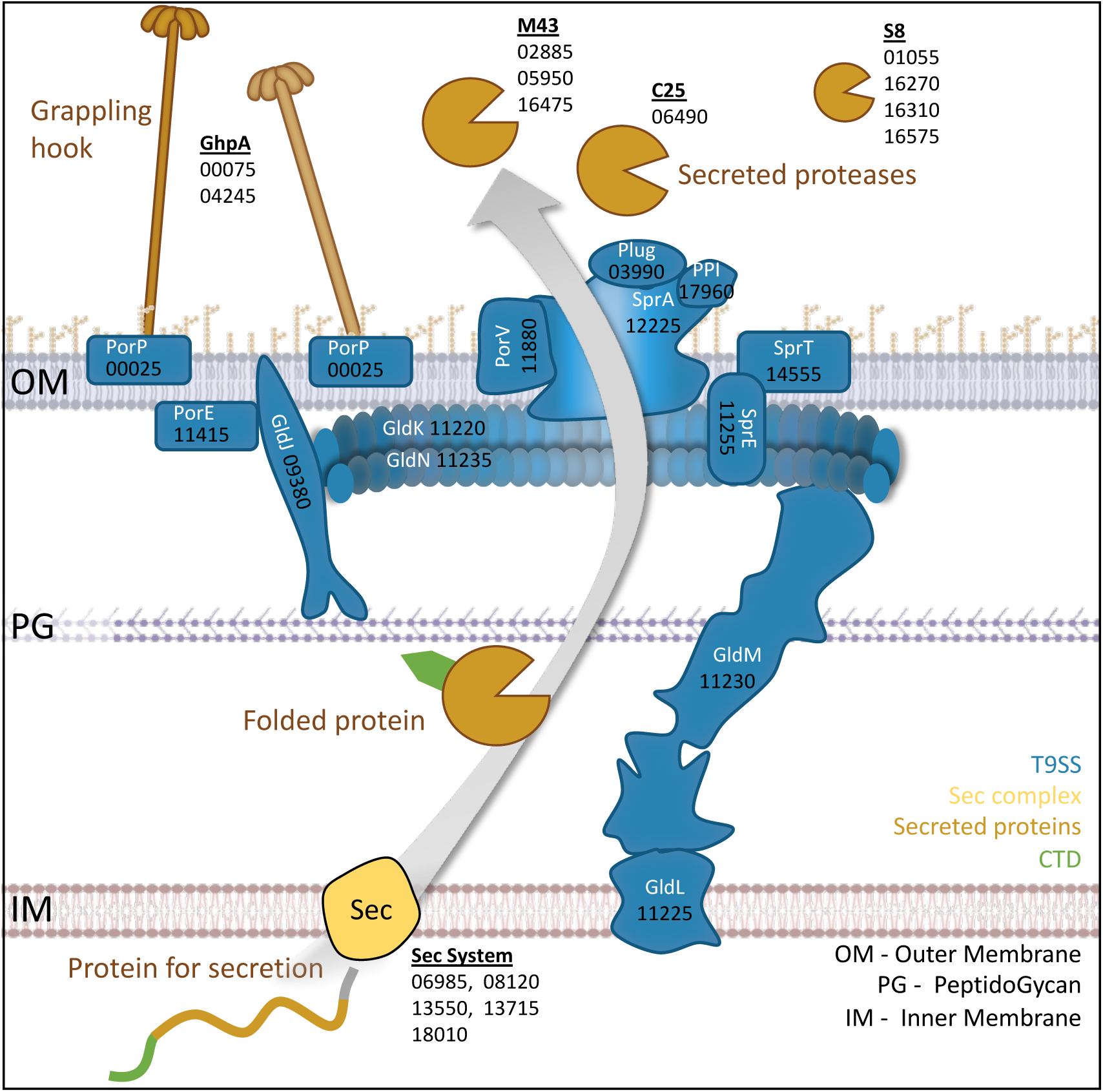
*E. algicida* Type IX Secretion System (T9SS) The *E. algicida* genome encodes for a complete T9SS (blue). Protein names (white), and protein IDs (black) are listed. Target proteins (ocher) for the T9SS are first recognized by the N-terminal signal peptide (gray) and secreted through inner membrane by the Sec system (yellow). The proteins fold in the intermembrane space, the C-Terminal Domain (CTD, green) is recognized by the T9SS, and the folded mature proteins are secreted out of the cell by the T9SS. Putative secreted proteases, M43 family Zn metalloprotease, C25 family cysteine peptidase and S8 family serine proteases are listed. Proteins with type B CTD remain anchored to the outer membrane by PorP proteins, for example the grappling hook protein A (GhpA) is presented. The full details of T9SS genes, and the putative secreted genes are presented in Table S6, and S7 respectively.

### *E. algicida* pathogenicity

The host range of *E. algicida* To15 pathogenicity was evaluated by adding the bacterium to a diverse set of 26 phytoplankton strains (Table 2, Table S8). We tested 10 *Thalassiosira* strains, along two other centric diatoms, seven pennate diatoms, and seven other strains of phytoplankton isolated from similar locations as *E. algicida* (*28*). Both *T. oceanica* (CCMP1005), and *T. pseudonana* (CCMP1335) were susceptible to *E. algicida* (Table 2), as well two strains of *T. diporocyclus* (CCMP3730, CCMP3731, Fig. S5) isolated from a similar transect as *E. algicida*. Each of these susceptible strains were axenic at the time of testing; two xenic *Thalassiosira* strains, *T. diporocyclus* CCMP3730, and *T. bioculata* CCMP3737 were also susceptible (Table 2). Two other xenic strains of *T. diporocyclus* (CCMP3729, CCMP3734, Table 2) also isolated from similar regions were resistant to *E. algicida* pathogenicity. Interestingly, the pennate diatom *Thalassiothrix* sp. (CCMP3718) was susceptible when axenic, but not when xenic (Table 2). None of the tested non-diatom strains appeared susceptible to *E. algicida*. Instead, the three tested *Pelagomonas calceolata* strains exhibited higher cell densities when co-cultured with *E. algicida* and a similar benefit was detected in one of the tested *Gephyrocapsa oceanica* strains (CCMP3746, Table 2). Overall, of the 26 tested phytoplankton strains, seven centric diatoms and two pennate diatoms were susceptible to *E. algicida*, including four of the eight strains initially used to enrich for algicidal filterable bacteria (marked with asterisks, Table 2); the tested non-diatom strains displayed either an enhanced growth response or no response to *E. algicida*.

**Table 2.**
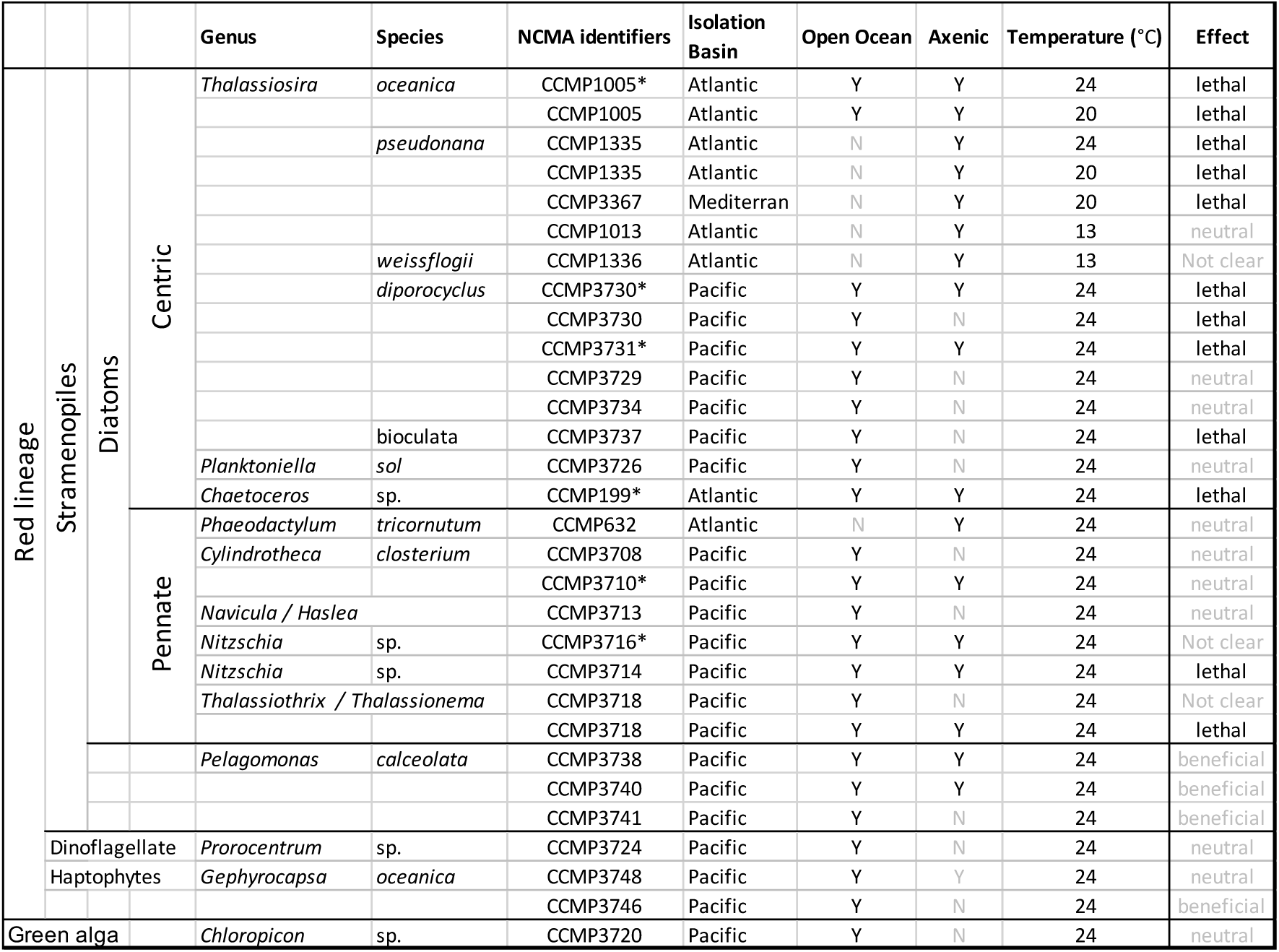
*E. algicida* host-range. Each tested host scientific name, CCMP culture collection ID, isolation basin, whether it is an open-ocean isolate, was the culture axenic when tested, and the temperature at which each strain was tested. The effect of *E. algicida* To15 on each strain is marked, strains that exhibited inconsistent results are marked as ‘not clear’. Asterisks mark strains that were part of the initial diatom mix used to enrich for algicidal bacteria.

We chose to focus on the interaction between *E. algicida* To15 and *T. oceanica* as it is a model pelagic diatom. *Ekhidna algicida* was co-cultured with *T. oceanica* and then filtered to obtain a diatom-free fraction of *E. algicida*; a new co-culture was initiated by adding these *E. algicidal* cells to an axenic *T. oceanica* culture at 1:10 v/v (about 20:1 *E. algicida* / *T. oceanica* cells). *Ekhidna algicida* cell abundance remained constant for the first 12 hours (6.7±1.3 million cells ml^-1^) of co-culture, increased by about 40% during day 1 (9.6±0.8 million cells ml^-1^), doubled over day 2 (17.4±3.3 million cells ml^-1^), and by days 4-7, reached final cell abundances of ∼40 million bacteria cells ml^-1^ (Fig. 3A). *Thalassiosira oceanica* co-cultured with *E. algicida* cells exhibited an immediate growth arrest (Fig. 3B) and after one day, a decrease in photosynthetic efficiency (Fv/Fm, Fig. 3C). The proportion of dead diatom cells in the co-cultures, as measured by percentage of Sytox positive cells, sharply increased during day 2, reaching 16.2±1.8% at 48 h and over 50.8±1.5% dead cells onward from 57 h (Fig. 3D). Monocultures of *T. oceanica* displayed a typical green-brown color after day 2 (54 h), whereas the *T. oceanica* + *E. algicida* co-cultures appeared clear (Fig. 3E). Examination of the co-culture using fluorescence microscopy revealed thin, elongated rod-shaped cells (length 2.6 - 5.3 µm, n = 16), surrounding the diatoms (Fig. 3F-G). Using transmission electron microscopy (TEM) we found that cells from the *T. oceanica* monoculture remained intact (Fig. 3H), whereas in co-culture with *E. algicida*, *T. oceanica* cells exhibited disruption of the frustule, disruption of the cell membrane, and expulsion of intercellular membranes and organelles (Fig. 3I-J).

**Fig. 3.**
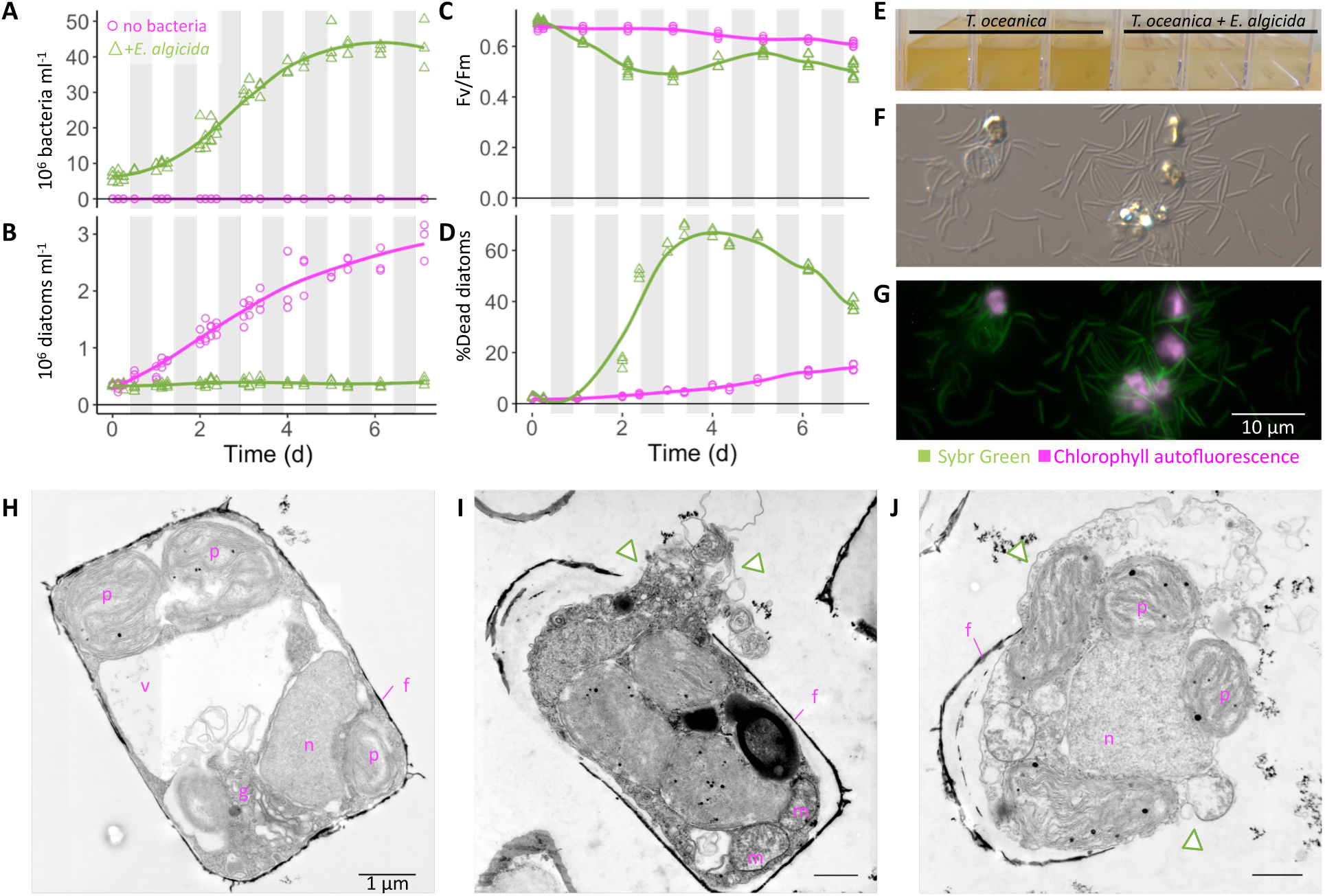
Interaction of *E. algicida* and *T. oceanica*. **A-D.** Co-culture of *T. oceanica* and *E. algicida* To15 added at t=0 (green triangles), and *T. oceanica* monocultures (magenta circles). Gray areas represent dark periods. **A.** Abundance over time of *E. algicida* cells, and similar-size fluorescent particles as detected by SYBR stain**. B.** Abundance over time of *T. oceanica* cells as detected by chlorophyll autofluorescence including cells with low chlorophyll autofluorescence. **C.** *T. oceanica* photosynthetic efficiency (Fv/Fm). **D.** Percentage of *T. oceanica* dead cells, measured as percentage of Sytox positive diatom cells. **E.** Image of the cultures taken at day 2, with triplicate flasks of *T. oceanica* monoculture (left) and *T. oceanica* + *E. algicida* (right). Phase (**F**) and fluorescence (**G**) microscopy images taken at day 7 of *T. oceanica* + *E. algicida* co-culture. Chlorophyll autofluorescence in magenta, the DNA stain SYBR-green in green. Scale bar 10 µm for both images. **H-J.** Electron microscopy images (TEM) of *T. oceanica* cells sampled at day 3. *T. oceanica* monoculture (**H**), and *T. oceanica* + *E. algicida* (**I-J**). In magenta letters: p – plastid, v – vacuole, n – nucleus, f – frustule, m – mitochondria, g – golgi. Expulsion of intracellular membranes and organelles out of the frustule are marked with green triangles. Scale-bars are 1 µm.

### Algicidal characteristic of *E. algicida* exudates

Given the pathogenic role of T9SS in other Bacteroidota species, we hypothesized that pathogenicity of *E. algicida* To15 would also involve proteins secreted via the T9SS (Fig. 2). The *E. algicida* To15 genome suggested a potential both for contact-dependent (by the grappling hook, and other attached proteins) and contact-independent pathogenicity (by the secreted protases). To distinguish between these two mechanisms, we designed experiments to test the pathogenicity of *E. algicida* exudates. *Thalassiosira oceanica* was grown as an axenic monoculture and in co-culture with *E. algicida*. After one week, the *T. oceanica* monoculture exhibited the characteristic green-brown color of a dense diatom culture, whereas the co-culture was essentially clear (Fig. 4A, scheme). Both the monocultures and the co-cultures were filtered through 0.2 µm pore-size filters to remove *T. oceanica* cells. Half of the < 0.2 µm fraction from the co-cultures were then filtered through 0.02 µm pore-size filters to achieve bacteria-free exudates (Fig. 4B, Fig. S6). Samples from the < 0.2 µm and the < 0.02 µm size fractions were added to fresh *T. oceanica* cultures at 0.4, 4 and 10% (v/v) (Fig. 4C). As a control, the exudates of *T. oceanica* monocultures were added at 10% (v/v) (Fig. 4B-C). Addition of *T. oceanica* monoculture exudates had no impact on the growth of fresh *T. oceanica* cultures (Fig. 4C-D, top panels). Addition of 4% or 10% (v/v) of the < 0.2 μm fraction, which contained *E. algicida* cells, prevented *T. oceanica* growth (Fig. 4C-D, middle panels, green triangles and squares); addition of 0.4% (v/v) of this same fraction allowed a slight increase in *T. oceanica* cell abundance between 2 to 5 days, followed by a subsequent decrease in cell abundance (Fig. 4C-D, middle panels, green circles). Addition of the bacteria-free < 0.02 μm fraction containing co-culture exudates to fresh *T. oceanica* resulted in a dose-dependent response: The 10% (v/v) exudates prevented any *T. oceanica* growth (Fig. 4C-D, blue triangles, bottom panels), whereas addition of 4% (v/v) exudates resulted in a lag phase of about 5 days before growth resumed, and addition of 0.4% (v/v) exudates resulted in a lag phase of about 2 days before growth resumed, with maximal growth rates of 1.58±0.15 and 0.74±0.09 respectively (Fig. 4C-D, blue squares and circles, bottom panel). These results indicate a contact-independent, dose-dependent pathogenicity of *E. algicida* exudates on *T. oceanica*. Live *E. algicida* cells propagate in co-culture and are lethal to *T. oceanica* regardless of the initial volume or concentration.

**Fig. 4.**
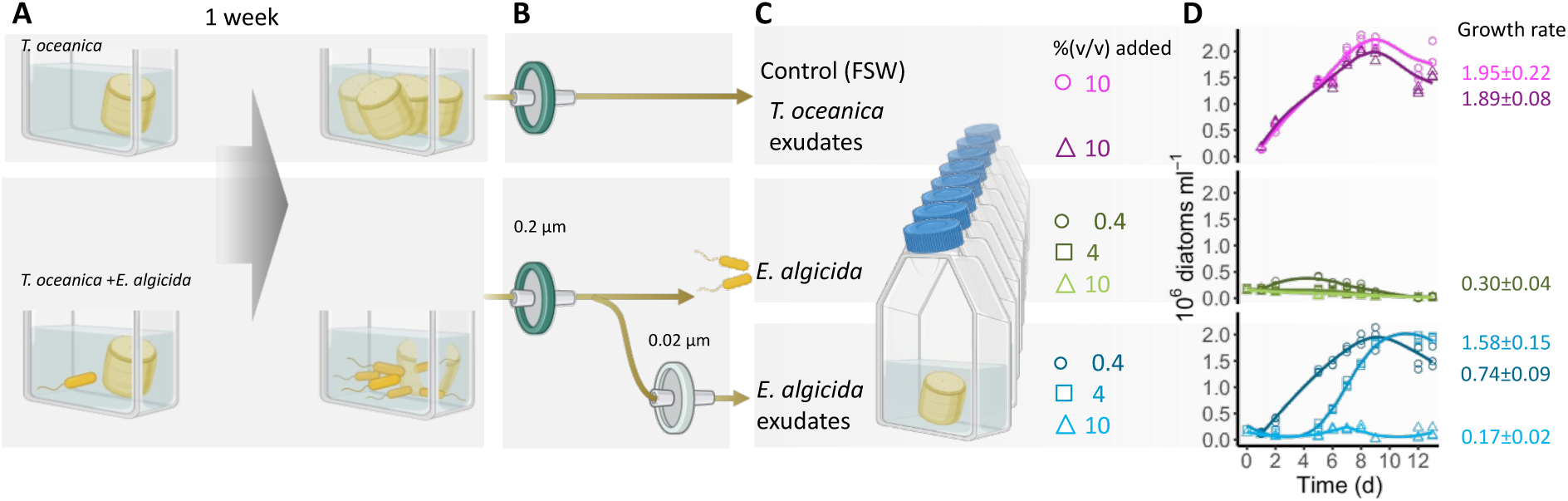
Exudates of *E. algicida* – *T. oceanica* coculture are algicidal to *T. oceanica*. **A-C.** Overview of experimental design. **A.** *T. oceanica* monocultures or co-cultures with *E. algicida* were grown for a week. **B.** Monocultures of *T. oceanica* were filtered through 0.2 µm pore-size filter to obtain *T. oceanica* exudates. Co-cultures of *T. oceanica* and *E. algicida*, were filtered through 0.2 µm pore-size filter, this fraction contained both the exudates and live *E. algicida* bacteria. An aliquot of the < 0.2 µm filtrate was filtered through a 0.02 µm pore-size filter to obtain bacteria-free co-culture exudates. **C.** Fresh *T. oceanica* cultures were treated with the above fractions added at different volumes to achieve 0.4%, 4%, and 10% (v/v). **D.** Growth curves of *T. oceanica* treated with filtered autoclaved seawater (FSW) (magenta circles) or *T. oceanica* exudates (dark purple triangles, top panel), with the < 0.2 µm fraction containing *E. algicida* and its exudates (green symbols, center panel), with the < 0.02 µm fraction of cell-free exudates (blues symbols, bottom panel). Maximum growth rates (division per day±SD) are indicated near the curves.

### *Ekhidna algicida* pathogenic switch in response to *T. oceanica*

We next asked whether the algicidal exudates produced by *E. algicida* were released constitutively or only upon exposure to *T. oceanica* or to *T. oceanica* secreted compounds. We grew *E. algicida* as both a monoculture in a glucose (3 M)-peptone (15 mg ml^-1^) media and in co-culture with *T. oceanica* (Fig. 5A, scheme). Addition of 10% (v/v) of the glucose-peptone media had no impact on *T. oceanica* growth yet still sustained *E. algicida* growth (Fig. S7). As observed in our previous experiments (Fig. 4), growth of *T. oceanica* was prevented by addition of the cell-free < 0.02 µm filtrate derived from the co-culture (Fig. 5A-B, blue triangles). In contrast, the cell-free < 0.02 µm filtrate from the *E. algicida* monoculture had no algicidal effect on *T. oceanica* (Fig. 5A, C, dark yellow triangles). Finally, we tested whether compounds released into the media by *T. oceanica* axenic monocultures could induce *E. algicida* to release pathogenic exudates. We first grew *E. algicida* in spent media of *T. oceanica* monoculture, supplemented with glucose (3 M) and peptone (15 mg ml^-1^) and then added the < 0.02 μm filtrate from this supplemented *E. algicida* monoculture to fresh *T. oceanica* cultures (Fig. 5A, D). These cell-free exudates had no effect on *T. oceanica* growth (Fig. 5D, dark green triangles). Together these results suggests that *E. algicida* produces and releases compounds into the media that are algicidal to *T. oceanica* upon interaction with *T. oceanica*, and that *E. algicida* monocultures do not produce such algicidal exudates.

**Fig. 5.**
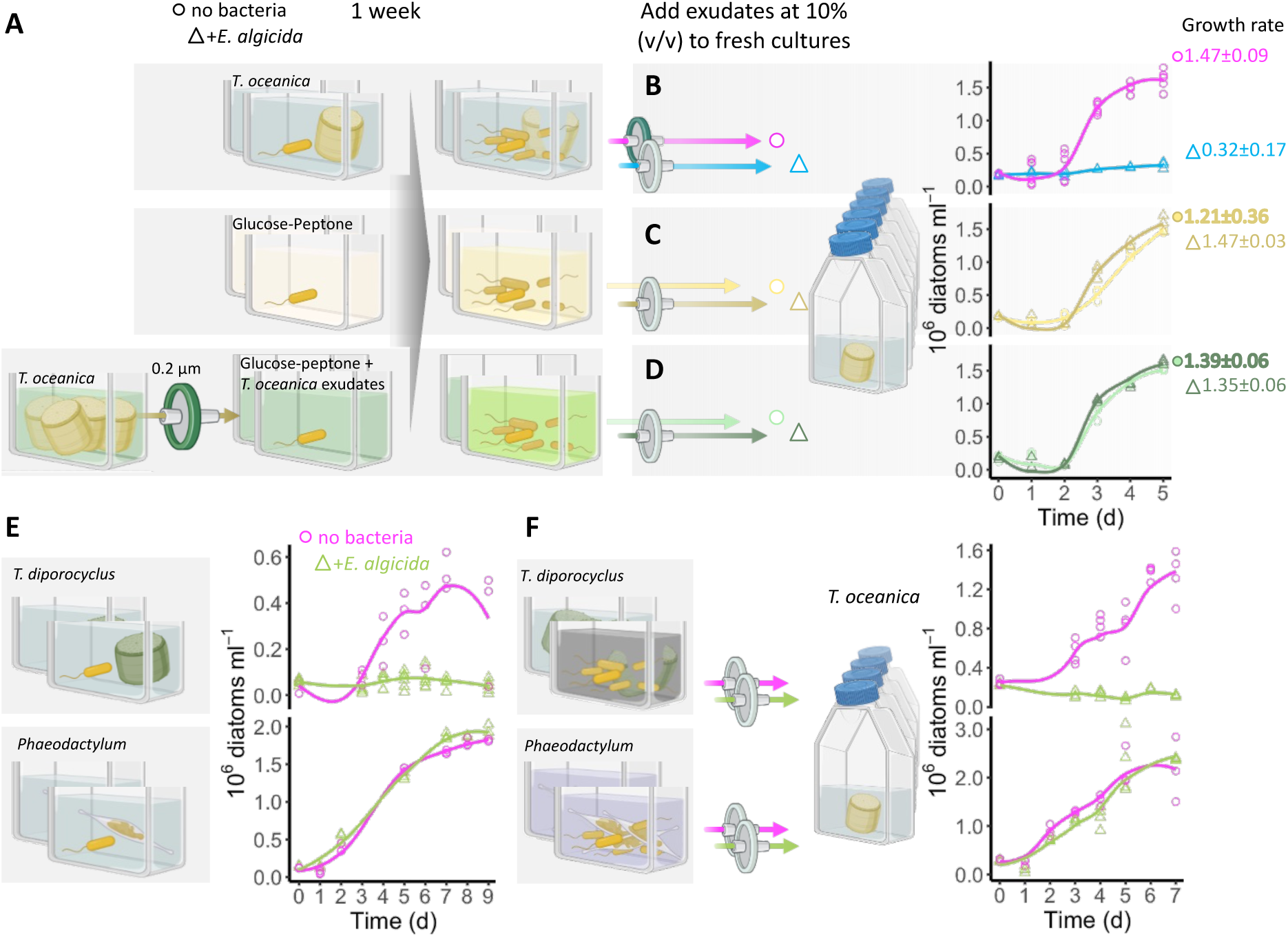
*Ekhidna algicida* pathogenicity switch. **A.** *Ekhidna algicida* cells were grown in co-culture with *T. oceanica* (top), as a monoculture in glucose-peptone media (middle), or as a monoculture in glucose-peptone media with exudates from *T. oceanica* monoculture (bottom). **B-D.** Each *E. algicida* culture was sequentially filtered through 0.2 µm and 0.02 µm pore-size filters to create bacteria-free exudates and added to fresh *T. oceanica* cultures (triangles). The different media were used as controls (circles). Maximum growth rates (division per day) are indicated near each growth curve. ***E*.** Diatom monocultures: top *T. diporocyclus* (CCMP3731), bottom *P. tricornutum* (CCMP632), were monitored with or without addition of *E. algicida* at t = 0, green triangles and magenta circles respectively. **F.** After about a week, the different diatom cultures with (green triangles) and without (magenta circles) *E. algicida* were sequentially filtered through 0.2 µm and 0.02 µm pore-size filters to achieve the cell-free exudates. The different exudates were added to fresh *T. oceanica* cultures that were monitored for a week.

We tested the lethality of *E. algicida* produced exudates under two additional co-culture scenarios: co-culture with the susceptible diatom *T. diporocyclus* or co-culture with the resistant diatom *Phaeodactylum tricornutum* (Table 2). We grew *T. diporocyclus* (CCMP3731) and *P. tricornutum* (CCMP632) as axenic monocultures (Fig. 5E, magenta circles), and in co-cultures with *E. algicida* (Fig. 5E, green triangles). As observed in previous experiments, *T. diporocyclus* did not grow in the presence of *E. algicida*, while *P. tricornutum* growth was not affected by *E. algicida* (Fig. 5E). In both co-cultures, *E. algicida* was able to grow as verified by OD_600_. We repeated the experiment, this time collecting the < 0.02 µm cell-free exudates from the diatom monocultures and the co-cultures after about a week. Each exudate was separately added at 10% (v/v) to fresh *T. oceanica* cultures (Fig. 5F). Exudates from the *T. diporocyclus*-*E. algicida* co-culture were algicidal toward *T. oceanica* (Fig. 5F, green triangles, top panel), whereas exudates from the *P. tricornutum*-*E. algicida* co-culture had no effect on *T. oceanica* (Fig. 5F, green triangles, bottom panel). These results suggest that both susceptible *T. oceanica* and *T. diporocyclus* can trigger a switch to pathogenicity by *E. algicida*. This same switch to pathogenicity is not apparent when co-cultured with a resistant diatom either because *P. tricornutum* does not elicit the switch or because this diatom is able to inactivate algicidal compounds within the exudate.

## Discussion

### Under-detection of filterable bacteria

Our isolation efforts show that algicidal *Ekhidna,* filterable through 0.2 μm filters, are present in the Pacific Ocean and can be enriched for by incubation with susceptible hosts. We suggest that inherent methodological biases have previously masked the presence of these bacteria. Notably, our new isolates were obtained from samples of several milliliters, whereas no or few *Ekhidna* ASVs were detected in samples of 1-4 L seawater concentrated on 0.2 µm pore-size filters in similar locations (Fig. 1A, compare isolates (green), with ASVs (purple and gray)). This study emphasizes the importance of use of different approaches to examine marine microbial diversity, as each approach carries its own inherent biases. While culture-independent methods are often biased by filter pore-size, culture-dependent methods are biased by the growth media, commonly selecting for bacteria that are fast-growing in rich media. However, *E. algicida* did not grow in MB, and exhibited a 2-day lag phase when grown in 50% MB (Fig. S7I). Selective cultivation of filterable bacteria, using diverse media types might expose a wider variety of open-ocean bacteria, whereas the use of bait organisms, as in this study, can be effective for isolation of pathogenic bacteria. *Ekhidna algicida* characterized here is ‘on the boundary’ of both methods: the bacterium can readily pass through a 0.2 µm pore-size filter, in co-culture and rich media the cells are elongated (Fig. 3F-G), such that a fraction of the cells can be trapped on 0.2 µm pore-size filters. Additionally, *E. algicida* failed to grow in standard rich media (MB). Accordingly, only one other *Ekhidna* species has been isolated (*E. lutea)*, despite identification of 76 *Ekhidna* MAGs in microbiome samples derived from the Pacific Ocean. Previous studies, dedicated to culturable filterable bacteria (concentrated on 0.1 µm pore-size filters) detected diverse filterable bacteria, including Bacterioroides, from diverse environments (*19*, *29*–*32*). In other studies, the < 0.2 µm fraction of seawater was flocculated with iron chloride to concentrate viruses, with diverse filterable bacteria also detected in this fraction, and a broad contribution of filterable bacteria to carbon fixation and cycling was suggested (*33*, *34*). Our results join these previous publications emphasizing the largely overlooked group of filterable, or ultra-small bacteria, that are present in the ‘dissolved’ or ‘viral fraction’ of seawater, in which their pathogenicity suggests a role in shaping community composition.

### The pathogenicity mechanism of *Ekhidna algicida*

The induction of *E. algicida* pathogenicity occurred only when co-cultured with susceptible diatoms (Fig. 5-6). Although we did not test whether this induction required cell-cell contact, once pathogenicity was induced, cell free *E. algicida* exudates were sufficient to kill *T. oceanica* through a contact-independent mechanism. Diatoms are encapsulated within a silica-based cell wall (frustule), embedded with proteins such as silacidins, silaffins, cingulins and frustulins (*35*, *36*) that enhance structural integrity of the frustule and inhibit silica dissolution. Electron microscopy images revealed breaks in the frustule, and expulsion of the diatom cell content in response to *E. algicida* (Fig. 3I-J). Bacterial extracellular enzymatic activity, including protease activity, can accelerate silica dissolution and damage the frustule (*37*, *38*). For example, AlpA1, an extracellular S8 serine protease secreted by the algicidal Bacteroidota *Kordia algicida*, is sufficient to kill diatoms (*39*). In another example a serine protease and metalloprotease produced by a marine *Pseudoalteromonas* exhibit algicidal activity against a diatom (*40*, *41*).

**Fig. 6.**
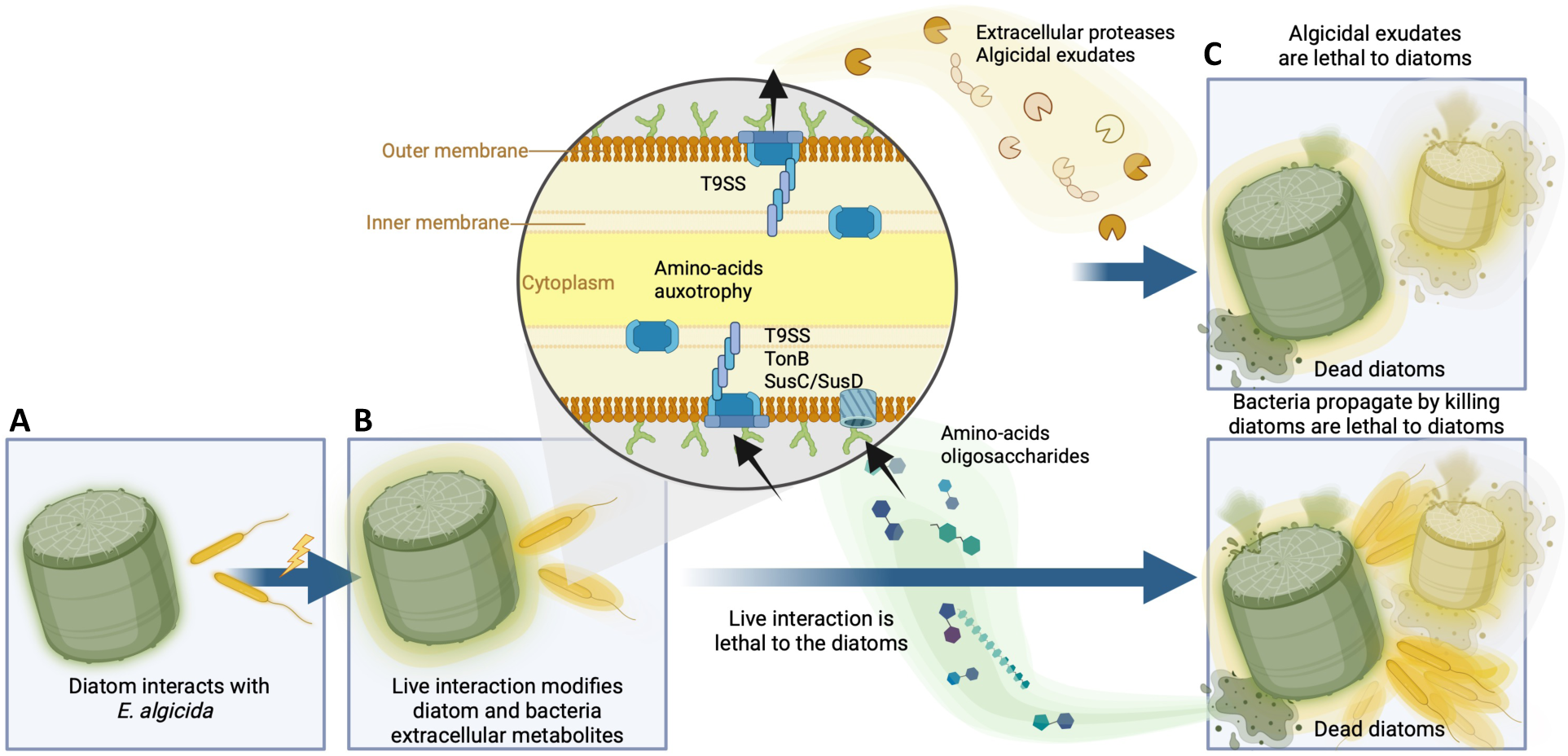
Conceptual model of *E. algicida* - diatom interactions. **A.** The diatom (green barrel), and *E. algicida* (yellow, elongated cells), are each surrounded by a faint halo representing steady-state exudates. **B.** Upon interaction, a pathogenic switch occurs and *E. algicida* modifies extracellular metabolites (the exudate), illustrated by more prominent haloes around the bacteria cells. Putative proteases and other algicidal metabolites are secreted through the T9SS. **C.** The *E. algicida* algicidal exudates produced upon live interactions are lethal to diatoms (top). Live interaction of *E. algicida* and diatoms is lethal to the diatoms and enables bacteria proliferation (bottom). Diatom lysis releases polysaccharides and amino acids that are essential to *E. algicida* growth. A combination of T9SS, TonB receptors, and SusC/SusD system facilitate uptake and degradation of large molecules such as oligosaccharides.

As prevalent in the Bacteroidota phylum, *E. algicida* encodes for T9SS (Fig. 2, Table S6), which relates to gliding, motility and the secretion of virulence factors (*21*, *42*, *43*). Accordingly, a search of *E. algicida* genome for proteins secreted by the T9SS revealed four S8 family serine proteases (Fig. 2, Table S7), that contribute to pathogenicity toward plants (*44*), and three M43 family Zn-metalloproteases (Fig. 2, Table S7) that participate in plant pathogenicity (*45*). Another putative algicidal protease is a C25 family cysteine protease (ACL8WY_06490), archetypes of this family are gingipains, notorious virulence factors, secreted through the T9SS that destroy periodontal tissue by *Porphyromonas gingivalis* (*43*, *46*). The T9SS can also secrete enzymes that break large molecules such as chitin and cellulose (*20*). Additional secreted glycoside hydrolase (ACL8WY_01805), heparin lyase (ACL8WY_14185), and potential oligosaccharide lyases such as xyloglucanase (ACL8WY_13760), cellulase (ACL8WY_14860), and alginate lyase (ACL8WY_16185), with overall 15 putative various sugar-hydrolases (Table S7) further suggest the ability to break the diatom cell wall and extracellular matrix, as well as initial metabolism of oligosaccharides. A combination of such extracellular carbohydrate-active enzymes with the SusC/SusD (Starch Utilization System) transporter pair function in large polysaccharide consumption in Bacteroidetes (*47*, *48*). We found seven SusC/SusD pairs in *E. algicida* genome (Table S9), suggesting polysaccharide utilization by *E. algicida* either from the diatom exudates, or when dying diatom cells lyse (Fig. 6).

While *E. algicida* exudates were sufficient to kill *T. oceanica* without contact in our experiments, it might not suffice in the open ocean. In our cultures, *E. algicida* grew to forty million cells per milliliter (Fig. 3A), which is orders of magnitude above the abundance of individual bacteria species in the open ocean. Therefore, in the dilute, oligotrophic pelagic environment, an additional contact-dependent mechanism may be required to maintain diatom-bacteria proximity and reach lethal concentrations of algicidal exudates around the diatom cell. Proteins secreted through the T9SS can be soluble or attached to the outer bacteria cell wall. Of the ten putative secreted proteins with type B CTD, seven were adjacent to *porP* genes (Table S7, Fig. S4B-C), an arrangement indicative of proteins that remain anchored via PorP to the outer cell membrane commonly involved in adhesion (*27*). We identified 25 extracellular proteins related to adhesion (Table S7), including the putative grappling hook protein A (GhpA) (ACL8WY_00030, 30.5% aa identity). The GhpA protein was characterized in *Aureispira* sp. (order Saprospirales, phylum Bacteroidota), in which the GhpA is secreted through the T9SS, and remains attached to the outer cell membrane, creating a grappling hook that is used it to catch and attach to prey bacteria (*49*). Other large (thousands of amino acids) extracellular adhesive proteins facilitate bacteria-prey and bacteria-host interaction, but are highly diverse and rapidly evolving, thus difficult to identify computationally (*50–52*). Of the 53 putative proteins secreted by the T9SS, 25 have a potential adhesive domain (Table S7). Furthermore, putative T9SS secreted proteins were significantly larger (average 1,300 aa) than expected by chance from *E. algicida* To15 genome (average 342 aa, *p* < 0.001, resampling test with 10,000 iterations). Using such large extracellular adhesive proteins, *E. algicida* can stay in the vicinity of the phytoplankton cell in the dilute pelagic environment for sufficient time to expiate its exudates and secrete enough algicidal exudates to kill and benefit from the death of the phytoplankton.

Some bacteria constantly produce algicidal exudates (*5*), while in other cases, the presence of a single algal metabolite can alter the bacteria exudates (*53–56*). Interestingly, we detected algicidal exudates produced from *E. algicida* upon live interaction with susceptible diatoms, but not when *E. algicida* was grown as monoculture in nutrient sufficient media, in supplemented diatoms spent media, or in live interaction with the non-susceptible diatom *P. tricornutum* (Fig. 5). It is yet to be determined if *P. tricornutum* does not trigger *E. algicida* pathogenicity or is actively resistant to *E. algicida*. For example, the diatom *Chaetoceros didymus* is resistant to the bacteria *Kordia algicida* by secreting a protease that degrades the algicidal protease of the bacteria (*57*). While bacteria isolated from rich coastal habitats can constantly secrete algicidal proteins (*5*, *15*), open ocean bacteria such as *Ekhidna* are likely frequently nutrient-limited, and might secrete algicidal exudates only when it is required and beneficial, such as upon interaction with susceptible diatoms. In general, the pathogenicity switch can be a response to specific metabolites produced upon interaction with susceptible phytoplankton, or from a lack of molecules or compounds required for bacterial growth, such as in low-nutrient environments (*14*, *58*). Whether this pathogenic shift is triggered by a general starvation response or a more specific signaling pathway remains to be determined.

### Ecological implications

We found algicidal *Ekhidna* bacteria in a wide area of the Pacific Ocean. *Ekhidna* had algicidal effects on open ocean *Chaetoceros* and *Thalassiosira* diatoms (Table 2), the two most abundant genera of diatoms in the oceans (*3*). Notably, *E. algicida* was algicidal toward three *Thalassiosira* strains, and two pennate diatoms species isolated from similar locations as the algicidal *Ekhidna* (Table 2), suggesting that such pathogenic interactions can occur in the open Pacific Ocean. The pathogenic interaction led to a disruption of the silica frustule (Fig. 3I-J), which can change the fate of the diatom’s organic carbon and silica (*59*). The ruptured diatom’s carbon content is likely to be recycled by *Ekhidna* as well as by other heterotrophic bacteria in the microbial loop rather than sink to the bottom of the ocean as dead intact diatoms.

### Conclusions

Microbial ecology is moving beyond describing the diversity of complex microbial communities and towards deciphering the functions of the plethora of uncharacterized microorganisms detected by molecular tools. In this context, the cultivation of prevalent but generally intractable microorganisms is important. *Ekhidna* bacteria are widespread throughout the Pacific Ocean and are algicidal to diverse pelagic diatoms. The algicidal mechanisms appear to be induced upon interaction with susceptible diatoms, and involve algicidal exudates, putatively including several proteases secreted by the T9SS. Characterization of pelagic diatom-bacteria interaction in the lab will facilitate further study and mechanistic understanding of such interactions in the Ocean.

### Protologue

*Ekhidna algicida* [al.gi.ci’da] L. fem. n. algi, alga; L. masc. n. suff. -cida, killer; N.L. masc. n. algicida, alga-killer referring to this organism killing microalgae.

Cells are yellow motile rods with length up to 5.2 µm (average 3.6±0.7 µm) and width of 0.4±0.1 µm (n=16). Cells can grow on 50% MB, seawater-peptone, or in co-culture with diatoms as the sole carbon and amino-acids source. The type strain, To15^T^, was isolated from the tropical Pacific Ocean at 16°N, 140°W at 15 m water depth. The DNA G+C content of the type strain is 40.38 mol% (determined from the genome sequence), and the genome is 3,967,350 bp in length. The type strain is To15^T^ (=ATCC TSD-518 = DSM 119715), GenBank accession number for the genome sequence is GCA_051379705.1.

## Material and Methods

### Phytoplankton cultures and maintenance

Phytoplankton cultures were grown in 12:12 hours light:dark cycles at 24 °C, or 16:8 hours light:dark cycles at 20 or 13 °C with light intensity of about 100 μmol photons·m^-2^·sec^-1^. Cultures were grown in filtered autoclaved Puget Sound seawater (FSW), supplemented with f/2, L1, or K nutrients (*60–62*). Details about each phytoplankton strain, whether the culture was axenic when tested, maintenance temperature and media are provided in Table S8. Note that the phytoplankton isolated from the Pacific open ocean were all isolated during the Gradients 4 and Gradients 5 cruises, as previously described (*28*). Other phytoplankton were obtained from the Provasoli-Guillard National Center for Culture of Marine Phytoplankton NCMA. Our isolated phytoplankton were deposited in NCMA and thus have NCMA identifiers (Table 2). We made some Pacific open ocean isolates axenic by antibiotic treatment as we previously described (*11*, *28*). Other axenic cultures were received axenic from NCMA. Presence of bacteria in all the axenic cultures was tested routinely by inoculation in MB, and by staining the cultures with SYBR green and visualization in epifluorescence microscopy.

### Algicidal bacteria isolation and culturing

Seawater samples were collected from various depths and locations during cruises in 2023 (Gradients 5, TN412), and 2024 (Early Career Chief Scientist Training Cruise, KM24-14) from several sources: the ship’s underway seawater system (UW), a trace-metal clean pump system (stayfish), and Niskin bottles, as indicated in Table S1. Seawater samples were filtered through 0.2 µm pore-size polyethersulfone (PES) syringe filters (13, or 25 mm diameter, Nalgene, ThermoFisher), or 0.2 µm pore-size polycarbonate filters (142 mm diameter, Sterlitech). *Ekhidna sp.* Tp1 was isolated from viral concentrate, generated by concentration of the 0.2 µm pore-size filtrate on a 30 kDa cut-off tangential flow filtration system as previously described (*63*). The sampling locations and collection dates of the isolates are indicated in Table 1 and Table S1. The 0.2 µm seawater filtrates were added to unialgal diatom cultures in 24 well plates or to a mixture of diatom cultures in 50 ml tissue-culture polystyrene flasks at about 10:1 v/v and maintained at 24 °C, 12:12 light:dark cycles. Approximately one week later, the cultures were filtered through 0.2 µm pore-size syringe filters, the filtrate was added to fresh unialgal diatom cultures at about 1:1 v/v, in 24 well plates, alongside sterile FSW as controls, and kept in the same growth conditions (Fig. S1A). A week later, clearer wells (indicative of pathogenicity) were filtered again through 0.2 µm pore-size syringe filters and added to fresh diatom cultures (same strain) at about 1:5 v/v in replicates in the first column of 24 well plates. From this column, 4 serial 1:10 dilutions were performed. After another week, the most diluted well that appeared clearer compared to the control was filtered through 0.2 µm syringe pore-size filters and added in a similar way to fresh diatom cultures. At the end of the cruise, the clearest wells were filtered through 0.2 µm pore-size syringe filters and kept in sterile micro-centrifuge tubes in the dark for 2 days during transport to the University of Washington laboratory. Upon return, the filtrates were added to fresh diatom cultures at about 1:50 ratio, and 4 serial dilutions were conducted as described above. After about a week, the most diluted clear wells were filtered and added to fresh diatom cultures as before, but this time with 8 replicates of 10 serial dilutions were conducted, up to 10^-10^ of the original. After about a week, filtrates from the most diluted clear wells were added to fresh diatom cultures in the same way. The most diluted clear wells of this treatment were considered clonal populations. The filtrates were kept in the dark at 4 °C and were all pathogenic toward *T. oceanica*. Bacterial strains were obtained from each filtrate tested in the lab. Individual algicidal bacteria strains were grown at room temperature in the dark in 2 ml in 24 well plates containing 50% MB (87.5% FSW, 2.5 g L^-1^ peptone, 0.5 g L^-1^ yeast extract). For cryopreservation, 800 µl of actively growing yellow cultures were added to 200 µl of sterile 100% glycerol, flash frozen in liquid nitrogen, and transferred to −80 °C. From this point, fresh cultures from glycerol stocks were used. The full details for each isolate sample location, date, depth, and the diatoms on which it was originally enriched are listed in Table S1.

### DNA extraction, sequencing, assembly and annotation

Clonal isolates of To15, Tp1, Tp2, Tp4, Tp6, To14, and To17, *Ekhidna* were grown in co-culture with *T. oceanica*, and 50 ml of each were filtered through a 0.2 µm, and the filtrate was used for extraction. All other isolates were grown as monocultures in 4 ml 50% MB. DNA was extracted using MasterPure^TM^ Complete DNA and RNA Purification (epicentre, MC89010), following the manufacturer’s instructions for Fluid Samples. Sanger sequencing of 16S rDNA was done by Genewiz (Azenta Life Sciences).

DNA from *E. algicida* To15 was sequenced using a combination of Oxford Nanopore long-reads and Illumina NovaSeq X Plus short reads. A circularized consensus contig of 3,967,350 bp was assembled within the same Geneious Prime® software package. The assembled genome and the raw read sequence files were uploaded to NCBI with the following identifiers: BioProject: PRJNA1214876, BioSample: SAMN46389320, genome assembly: ASM5137970v1, Genebank: GCA_051379705.1, genome raw reads, SRA: PRJNA1214876. The genome was automatically annotated by NCBI Prokaryotic Genomes Annotation Pipeline (*64*). ANI analysis was done using JSpecies Web Server (JSpeciesWS) (*65*). Amino acids auxotrophies were predicted in GapMind (*66*, *67*).

Putative proteins secreted by the T9SS were identified by the hmm search, with an e value of 0.001 and hmm profiles of the two characterized types of C-Terminal Domains (CTD) for T9SS: type A CTD (PF18962, TIGR04183), and the type B CTD (PF13585, TIGR04131) (*68*, *69*). Detection of the N-terminal Sec signal peptide was done using Phobius (http://phobius.sbc.su.se/) (*70*). The putative secreted proteins were searched using HHpred (*71*).

All sequence alignments were done using Multiple Alignment using Fast Fourier Transform (MAFFT) version 7.313 (*72*, *73*); parameters: localpair, 100 iterations. A maximum-likelihood 16S phylogenetic tree was built using RAxML version 8.2.4 (*74*); parameters: -f a -x 75601 -p 301424 -# 100 -m GTRGAMMAI. Genomes tree (parameters:-f I -m GTRCAT) was based on 100 single copy genes (Table S3), and was done directly in BV-BRC (*75*). All tree visualizations were performed in the Interactive Tree of Life version 7 (https://itol.embl.de/; (*76*)).

### Pathogenicity experiments, using live bacteria or their exudates

*Ekhidna algicida* bacteria were added to axenic exponential cultures of *T. oceanica* grown at 24 °C. For the experiments presented in Fig. 3, < 0.2 µm filtrate of 1 week infection of *T. oceanica* with *E. algicida* was added in 1:10 dilution. In Fig. 4 and Fig. 5, *E. algicida* was grown in the various media types, co-culture with diatoms were done in the diatom growth conditions, and *E. algicida* growth in monoculture was at room temperature (∼ 24 °C) in the dark. Cell-free exudates were achieved by serially filtering through a 0.2 µm pore-size PES syringe filters, and then through a 0.02 µm pore-size inorganic membrane filter (Anopore, Whatman). Replicate cell-free filtrates were tested by 1:10 dilution into fresh 50% MB in multiwell plates. Plates were kept at room temperature and monitored for visible bacteria growth for at least 2 weeks (example in Fig. S6).

### Cell counts and physiology measurements

*T. oceanica* and *E. algicida* counts presented in Fig. 3A-B were run on a BD Influx flow cytometer (BD, Franklin Lakes, NJ, USA) (488 nm laser; emission 692±40, 530±40 nm) with at least 10,000 cells analyzed per sample. Cells were gated using the FCSplankton package in R (https://github.com/fribalet/FCSplankton). Two ml samples were incubated for 20 min with 2% v/v glutaraldehyde, flash frozen in liquid nitrogen, and stored at −80 °C prior to analysis. *E. algicida* were counted after staining with SYBR Green I (1:10000) for 20 minutes. For all other experiments (Figs. 4-5), samples of 150 µl were taken and measured immediately using Guava easyCyte 11HT Benchtop Flow Cytometer, excitation by 488 nm laser. Phytoplankton cells were detected by chlorophyl autofluorescence (680±30 nm) and forward scatter. *T. oceanica* cell death was determined by positive Sytox Green (Invitrogen) staining, used at a final concentration of 1 µM. Samples were incubated in the dark for 30 minutes prior to measurement. Positive staining (525±30 nm) was determined according to untreated stained cells and treated unstained cells.

Photosynthetic efficiency: Photochemical yield of photosystem II (Fv/Fm) was determined with a Phyto-PAM fluorometer (Heinz Walz GmbH, Effeltrich, Germany) using 15 min dark-adapted cells. Triplicate samples were measured as 1 ml in 1 cm cuvette, and each sample was measured 3 times with 20 sec intervals.

Light microscopy: Live cells were stained with SYBR Green I (1:10000) for 20 minutes. Cells were visualized using Nikon Eclipse 80i upright microscope (Nikon, Tokyo, Japan) equipped with differential interference contrast, 100x magnification, a blue-excitation, 515 nm, long-pass emission filter set (11001v2; Chroma Corp., Rockingham, VT, USA), and a MicroPublisher 3.3 RTV color CCD camera (QImaging, Burnaby, BC, Canada). Chlorophyll autofluorescence and brightfield images of live cells were taken at 20x magnification (Leica DMi8 microscope).

### Electron microscopy

At 2 days post infection, 75 ml of *T. oceanica* monoculture, and 75 ml *T. oceanica* + *E. algicida* co-culture were centrifuged for 10 min at 4000 rpm, the supernatant was removed, leaving 1.5 ml of sample. The resulting sample pellets were centrifuged at 7500 rpm for 1-2 minutes to further concentrate the cells. An equal amount of (primary) fixative consisting of 3% gluteraldehyde/2% paraformaldehyde buffered with a final 0.1 M sodium cacodylate buffer was added to the samples. Samples were resuspended and fixed at room temperature for 2 hours. Following 2 hours of fixation, samples were centrifuged for 1-2 minutes, and fixative was rinsed with 0.1 M sodium cacodylate buffer 4 × 10 minutes each. A post fixation was done with 1% osmium tetroxide (OsO_4_) buffered with 0.1 M of the same buffer for 1 hour. Following the OsO_4_ fixation, samples were centrifuged and rinsed in the same manner as with the primary fixative. Dehydration was done with a graduated series of 50%, 70%, 90%, 100% (twice) ethanol and 100% acetone (2 X) 10 minutes each.

Infiltration and embedding for TEM: Following the last 100% acetone dehydration step, the solvent was changed to 50:50 acetone:resin (Mollenhauer II), samples were resuspended, caps were secured and sealed with parafilm and put on a slow-moving rotator at room temperature for 24 hours. After the 24 hours of infiltration, the samples were again pelleted by centrifugation and were then returned to the rotator for about 2 hours, with caps removed to evaporate acetone. After 2 hours the pellet was carefully removed from the tubes and placed into gelatin capsules of fresh 100% resin Mollenhauer II and put into a vacuum chamber (under light vacuum) for 2 hours. After 2 hours in the vacuum chamber, the pellets were divided into several samples and placed in fresh 100% Mollenhauer II resin in flat molds and polymerized in a 60° C oven overnight.

Microtoming and TEM imaging: Embedded blocks were trimmed with a razor blade and microtomed using a 30 degree diamond knife (Diatome, PA, USA) with a Leica UC-6 ultramicrotome. Sections were cut at 80 nm and collected on 150 mesh carbon only TEM grids (Electron Microscopy Sciences, PA, USA). Grids were stained using a standard positive staining protocol with 2% uranyl acetate and lead citrate. Grids were imaged with an FEI (now ThermoFisher Scientific) Tecnai F20 at 200 kV utilizing the HAADF (s)TEM detector for optimal contrast. Selected images were minimally processed by Inkscape: colors were inverted and the images cropped.

### Collection, sequencing and detection of *Ekhidna* in ASVs

Four datasets, each collected in a different year along one of two transects, were sampled and analyzed in the North Pacific. Gradients 1-3 were collected along a latitudinal transect during three separate cruises in 2016 (Gradients 1; KOK1606), 2017 (Gradients 2; MGL1704), and 2019 (Gradients 3; KM1906), as described by Key et al. (2025) (*77*). The Gradients 4 cruise (TN397) was collected in 2021 along a separate transect, as described by Jones-Kellett et al. (2024) (*78*). Gradients 1–3 ASV samples were collected using 0.2 μm pore-size Supor membranes (Pall Corporation) and 3.0 μm pore-size polyethersulfone membranes (Sterlitech) and amplified using the 515F/806R primer set (*79–81*), while Gradient 4 ASV samples were collected on 0.22 μm pore-size filters and amplified using the 515Y/926R primer set (*78*). The sequences are available in NCBI under accession number PRJNA1079727.

Type A *Ekhidna* 16S rDNA was used as a blast query, we consider sequences with up to 4 mismatches (>97% identity) with any of our *Ekhidna* isolates as detected algicidal *Ekhidna*. Type A, and ‘*Ekhidna*’ 16S rDNA were searched in GLOSSary (*82*), restricting results to chimeraFree contigs over 900 bp.

## Acknowledgment

We thank the SCOPE-Gradients team for assisting with sample collection, sharing data, and providing helpful feedback that improved this work, and the scientists, captain, and crew of the R/V Kilo Moana and R/V Thomas G. Thompson. K, and specifically Angelicque E. White and Matthew Church, chief scientists on the KN24-14 cruise. We thank Kelsy Cain for flow cytometry analysis, Wenyu Shi and Celeste Castaneda-Lopez for testing the host range. We thank Debbie Lindell and Laure Arsenieff for advice on how to isolate filterable pathogens and Sigitas Sulcius for providing the < 0.2 µm concentrate sample, from which Tp1 was isolated. We thank Robert Morris and Michael Sadler for assistance with nanopore sequencing, and valuable feedback.

## Author contributions

Conceptualization: SGvC, EVA

Methodology: SGvC, EL, JM Formal analysis: VI

Investigation: SGvC, RM, MJS, JM, VI, AEJK, JM, RSK

Visualization: SGvC, EL

Supervision: EVA, BPD, JF

Writing—original draft: SGvC, SNC, EVA

Writing—review & editing: SGvC, SNC, EL, VI, RM, MJS, AEJK, JM, RSK, JF, BD, EVA

## Data and materials availability

All data needed to evaluate the conclusions in the paper are available in the paper and/or the Supplementary Material. *E. algicida* To15 is available from the American Type Culture Collection (=ATCC TSD-518) and the German Collection of Microorganisms and Cell Cultures (=DSM 119715). Algal cultures are available from the Provasoli-Guillard National Center for Culture of Marine Phytoplankton (NCMA), with the identifiers indicated in Table 2.

