## Supplementary Figures for "Induced pathogenicity toward open-ocean diatoms by a newly isolated filterable bacterium *Ekhidna algicida* sp. nov."

### Supplementary Figures and Legends

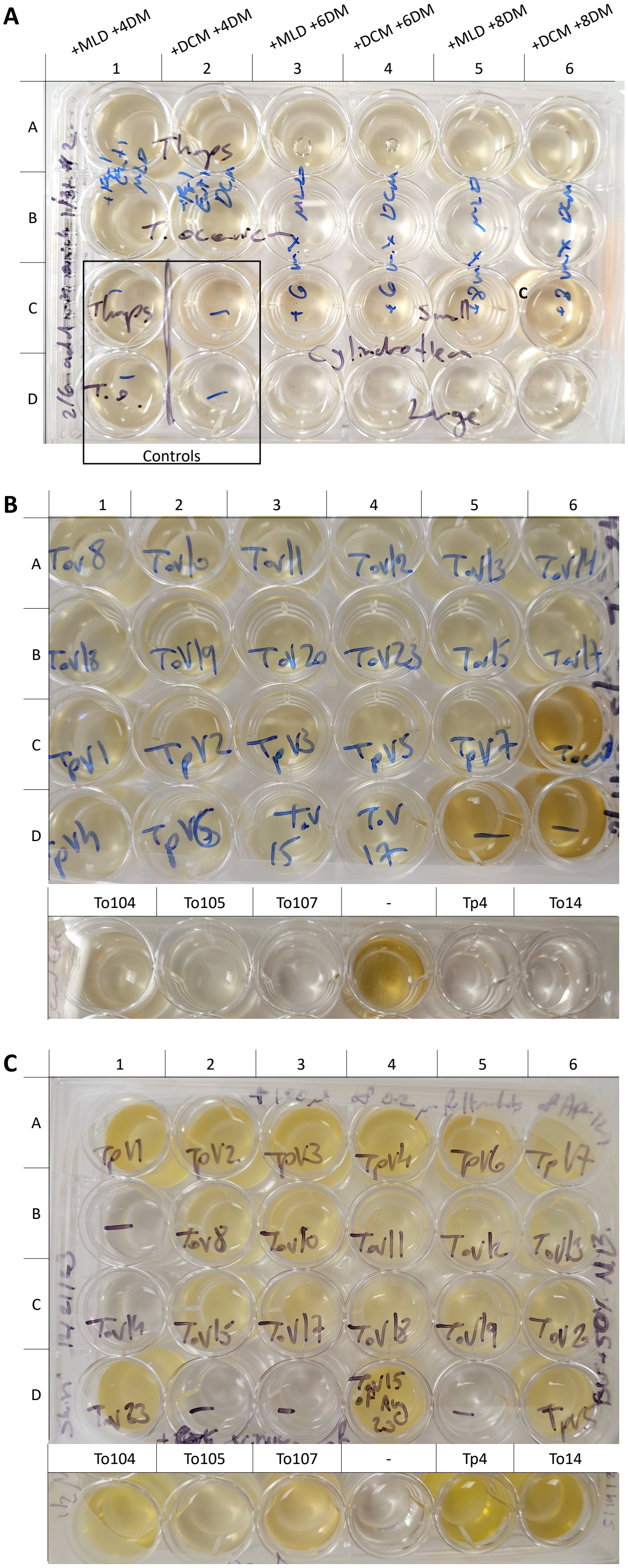


#### Figure S1 Isolation of filterable algicidal bacteria

**A.** An example of enrichment and initial isolation of filterable algicidal bacteria, by addition of < 0.2 µm filtrates to diatom cultures. Samples from 12.21°N, 140°W, 45 m (the mixed layer depth - MLD), or from 95 m (the deep chlorophyll maximum - DCM), were filtered through a 0.2 µm filter, and the filtrate was added to mixture of different diatoms mixtures (DM): 4DM, 6DM or 8DM (see Table S1). After 6 days the different DMs were filtered through 0.2 µm filter, and 250 µl of the filtrates were added to wells containing 2 ml of *T. pseudonana* CCMP1335 (A1-6, C1), *T. oceanica* CCMP1005 (B1-6, D1), *Cylindrotheca closterium* CCMP3709 (C2-C6) or *C. closterium* CCMP3710 (D2-D6). Control wells (C1, C2, D1, D2) were treated with 250 µl of sterile autoclaved filtered seawater (FSW). Image was taken after 4 days and showed reduced growth of *T. oceanica* in wells B3-6 relative to wells B1, B2, and D1. Strain To20 was isolated from well B6, and strain To23 was isolated from well B3. **B.** Bacteria effect on *T. oceanica* cultures. One hundred microliter of < 0.2 µm filtrate of each of the 22 isolates was added to 2 ml of fresh *T. oceanica* cultures. The plates were imaged after a week. Wells without added filtrate are marked by a dash and exhibit the green-brownish color of a dense diatom culture (wells D5, D6- top plate, well 4 bottom plate). Control with < 0.2 µm filtrate of axenic *T. oceanica* culture is in well C6. **C.** Bacteria monocultures. One hundred microliter of < 0.2 µm filtrate of each isolate was added to 2 ml of 50% MB. Control wells (no added filtrate) are marked a hyphen (wells B1, D2, D3, D5- top plate, well 4 – bottom plate). The plate was imaged after 10 days, showing that all the bacteria isolates can visibly grow in 50% MB, mostly exhibiting bright yellow color.

### Figure S2 *Ekhidna* 16S rDNA

**A**

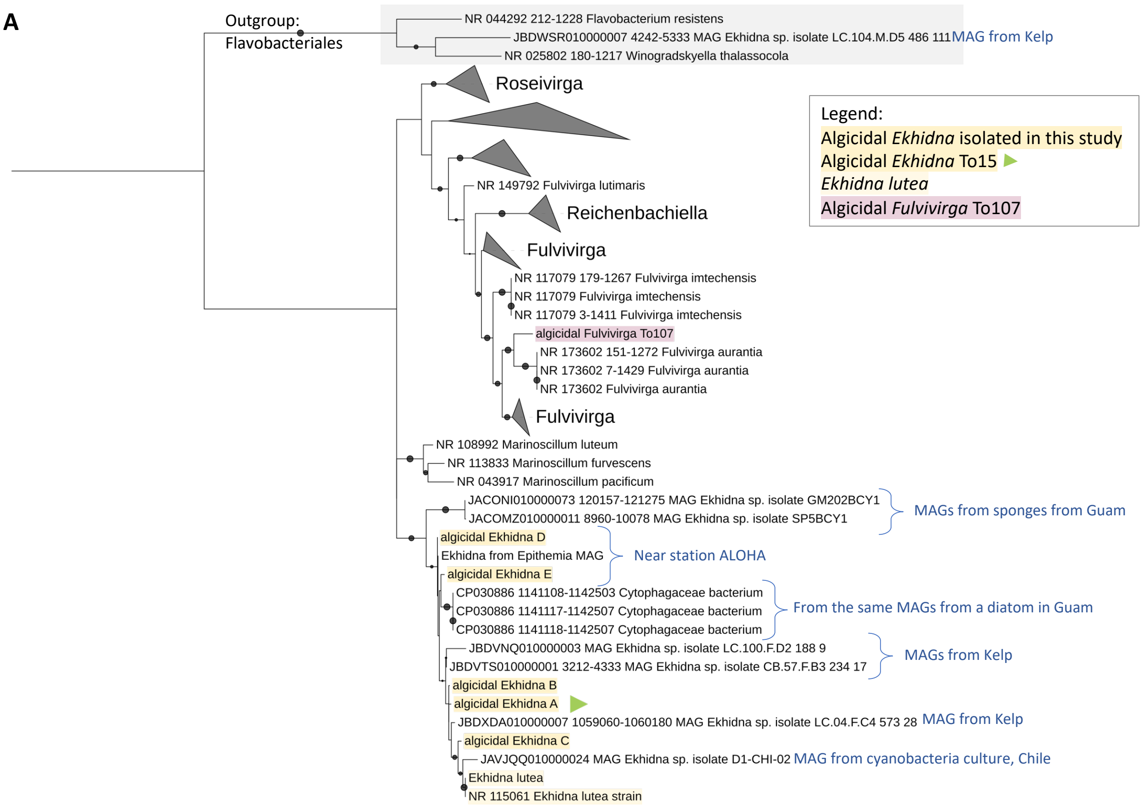


**B**

```
E. algicida A  1     ......G......................................................................T...........................TA...........................................
Ekhidna B      1     T.....G..............................................G.......................T...........................TA...........................................
Ekhidna E      1     ......G....T....................A....................G...................................................CA...........................................
Ekhidna D      1     ......G..............................................G...................................................GA...........................................
Ekhidna C      1     ..T..........................................................................T...........................GAT..........................................
E. lutea       1     CTCCATAGTATCACAGATAGGCATCTTGAAGTGATTAAAGATTTATCGGTATTAGATGAGTATGCGTCTGATTAGTTAGTTGGTGAGGTAACGGCTCACCAAGACAGCGATCAGTAGGGGTTCTGAGAGGATGATCCCCCACACTGGCAC

E. algicida A  151   ..........................................................................................................................T............G..T....A..T...
Ekhidna B      151   ..........................................................................................................................T............G..T....A..T...
Ekhidna E      151   ...........................................................A..............................................................T............G..T....A..CA..
Ekhidna D      151   ..........................................................................................................................T............GC.T....A.GCA..
Ekhidna C      151   ..........................................................................................................................T...........................
E. lutea       151   TGAGATACGGGCCAGACTCCTACGGGAGGCAGCAGTAGGGAATATTGGTCAATGGGCGAGAGCCTGAACCAGCCATGCCGCGTGCAGGATGACGGCCTTCTGGGTTGTAAACTGCTTTTATACGGGAAGAAAAAGCTCATGCGTGAGCAA

E. algicida A  301   ...........A..T.......................................................................................................................................
Ekhidna B      301   ...........A..T.......................................................................................................................................
Ekhidna E      301   ...........A..T.................................................................................................C......................G..............
Ekhidna D      301   ...........A..T.................................................................................................C......................G..............
Ekhidna C      301   ...........A..T........................................................................................................................G..............
E. lutea       301   TTGCCGGTACCGTACGAATAAGCACCGGCTAACTCCGTGCCAGCAGCCGCGGTAATACGGAGGGTGCAAGCGTTGTCCGGATTTATTGGGTTTAAAGGGTACGTAGGCGGATTTTTAAGTCCGTGGTGAAAGCCTACAGCTTAACTGTAG

E. algicida A  451   ......................................................................................................................................................
Ekhidna B      451   ......................................................................................................................................................
Ekhidna E      451   ...................G..................................................................................................................................
Ekhidna D      451   ...................G..........................................................C.......................................................................
Ekhidna C      451   ..................G...................................................................................................................................
E. lutea       451   AACTGCCATGGATACTGGAAATCTTGAATTCAGTTGAGGTAAGCGGAATTTATGATGTAGCGGTGAAATGCATAGATATCATAAAGAACACCTATTGCGAAGGCAGCTTGCTGGACTTGAATTGACGCTGAGGTACGAAAGCGTGGGGAG

E. algicida A  601   ......................................................................................................................................................
Ekhidna B      601   ......................................................................................................................................................
Ekhidna E      601   ......................................................................................................................................................
Ekhidna D      601   ......................................................................................................................................................
Ekhidna C      601   ......................................................................................................................................................
E. lutea       601   CGAACAGGATTAGATACCCTGGTAGTCCACGCCGTAAACGATGATTACTCGCTGCTGGCGATACACTGTCAGTGGCCAAGCGAAAGCGTTAAGTAATCCACCTGGGGAGTACGCTCGCAAGAGTGAAACTCAAAGGAATTGACGGGGGTC

E. algicida A  751   ......................................................................................................................................................
Ekhidna B      751   ......................................................................................................................................................
Ekhidna E      751   ......................................................................................................................................................
Ekhidna D      751   ......................................................................................................................................................
Ekhidna C      751   ......................................................................................................................................................
E. lutea       751   CGCACAAGCGGTGGAGCATGTGGTTTAATTCGATGATACGCGAGGAACCTTACCTGGGCTAGAATGTGAGCGCAATCTCTAGAGATAGAGAGTTCTTCGGACGTGAAACAAGGTGCTGCATGGCTGTCGTCAGCTCGTGCCGTGAGGTGT

E. algicida A  901   ....................................................................T...................................C.............................................
Ekhidna B      901   ....................................................................T...................................C.............................................
Ekhidna E      901   ......................................................................................................................................................
Ekhidna D      901   ......................................................................................................................................................
Ekhidna C      901   ........................................................................................................C.............................................
E. lutea       901   TGGGTTAAGTCCCGCAACGAGCGCAACCCCTACTGTTAGTTGCCAGCATGTAATGATGGGGACTCTAACAGGACTGCCTACGCAAGTAGAGAGGAAGGAGGGGATGACGTCAAGTCATCATGGCCCTTACGCCCAGGGCTACACACGTGC

E. algicida A  1051  ................................CA......................................
Ekhidna B      1051  ........................................................................
Ekhidna E      1051  ................................CA......................................
Ekhidna D      1051  ........................................................................
Ekhidna C      1051  ........................................................................
E. lutea       1051  TACAATGGAGGGTACAGAGGGTCGCTAGCCAGTGATGGTATGCCAATCTCAAAAAGCCCTTCTCAGTTCGGA
```

**A.** Phylogenetic tree of *Ekhidna* and Cytophagales 16S rDNA sequences derived from isolates and publicly available metagenome assembled genomes (MAGs). Black circles indicate bootstraps > 70. The outgroup, Flavobacteriales, is marked with gray background. Triangles indicate collapsed groups. *Ekhidna* isolates are marked with light yellow background, *E. algicida* To15 16S rDNA (*E. algicida* A) is marked with a green triangle. Algicidal *Fulvivirga* To107 is marked with pink background. Samples origin of *Ekhidna* 16S rDNA derived from MAGs are indicated. **B.** *Ekhidna* 16S rDNA sequence aligned to *E. lutea*. Nucleotides identical to *E. lutea* are marked with dots.

Figure S3 A 100 genes codon tree of *E. algicida* To15, *E. lutea* and *Ekhidna* MAGs

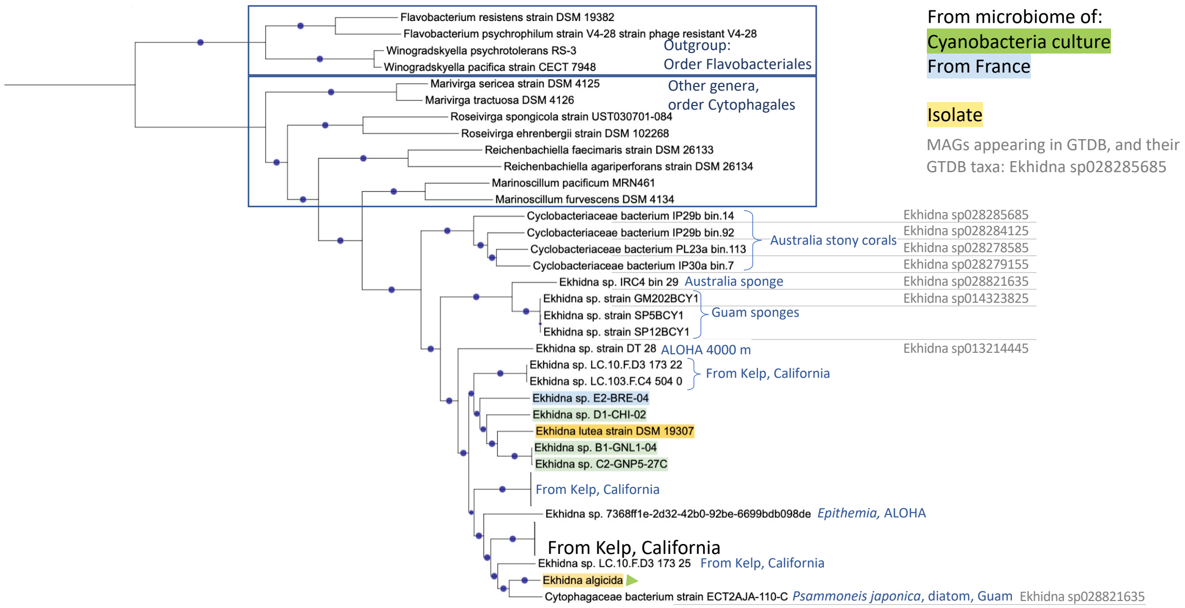


A maximum likelihood phylogenetic tree derived from 100 single copy genes of *Ekhidna* genomes. Black circles indicate bootstrap > 70. *Ekhidna* isolates are highlighted in yellow background, *E. algicida* To15 is marked with a green triangle. Sample origins of MAGs are indicated in blue. GTDB species IDs are indicated in gray when available.

### Figure S4 Alignment of the CTDs of putative proteins secreted by T9SS

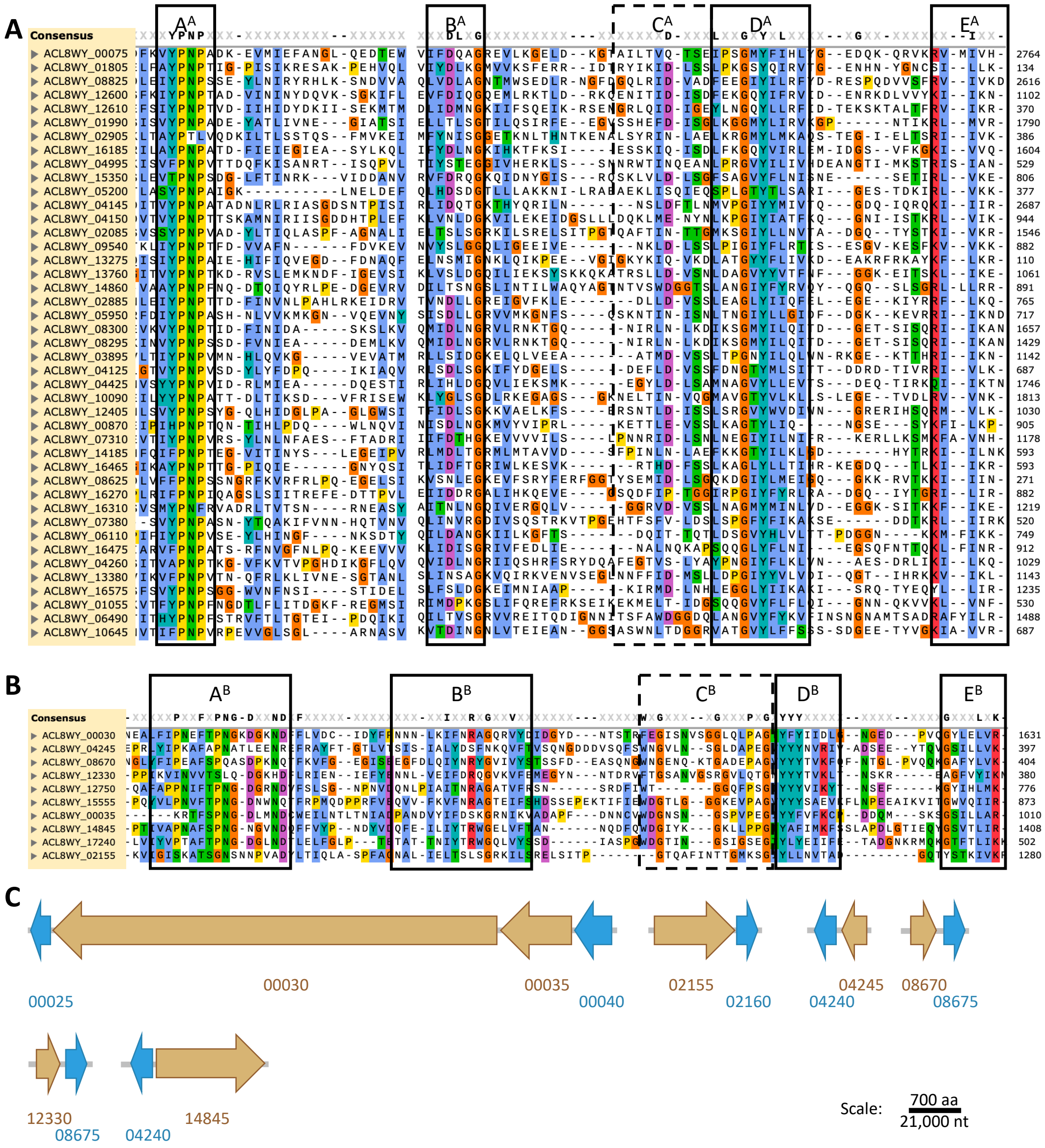


Alignment of the type A CTD (**A**), and type B CTD (**B**) of *E. algicida* To15 putative proteins secreted by T9SS. Consensus is shown for >50% conservation, and aa are colored by Clustal X scheme. Protein IDs precede the sequences. Structural conserved regions are marked by black boxes according to Paillat et al. (*1*). **C**. The genomic architecture of putative proteins secreted with type B CTD (ocher), that are adjacent to genes encoding to PorP proteins (blue). The translation directions are marked by the arrow directions.

### Figure S5 Examples of diatoms susceptible to *E. algicida* To15

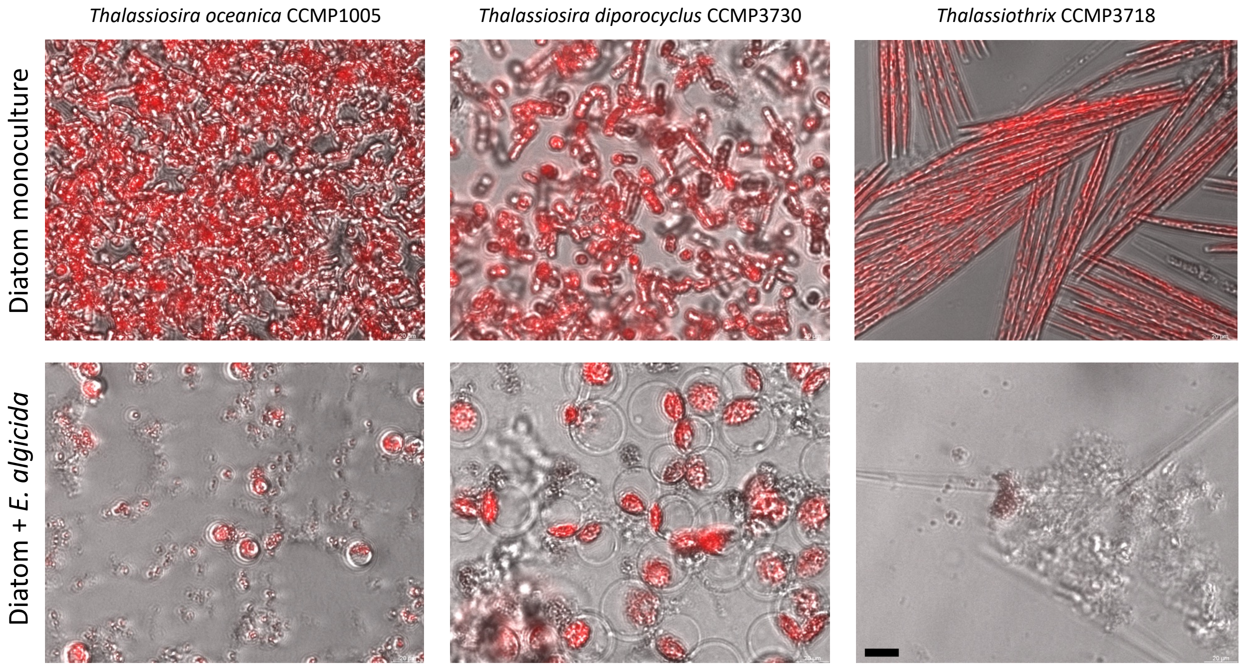


Overlay of light and fluorescence microscopy images of dense cultures of axenic diatoms (top), and the same diatoms in co-culture with *E. algicida* To15 (bottom). Chlorophyll autofluorescence is presented in red. Magnification is the same for all images, the scale bar is 20 µm.

### Figure S6 *Ekhidna algicida* cells pass through 0.2 µm and not through 0.02 µm pore-size filters

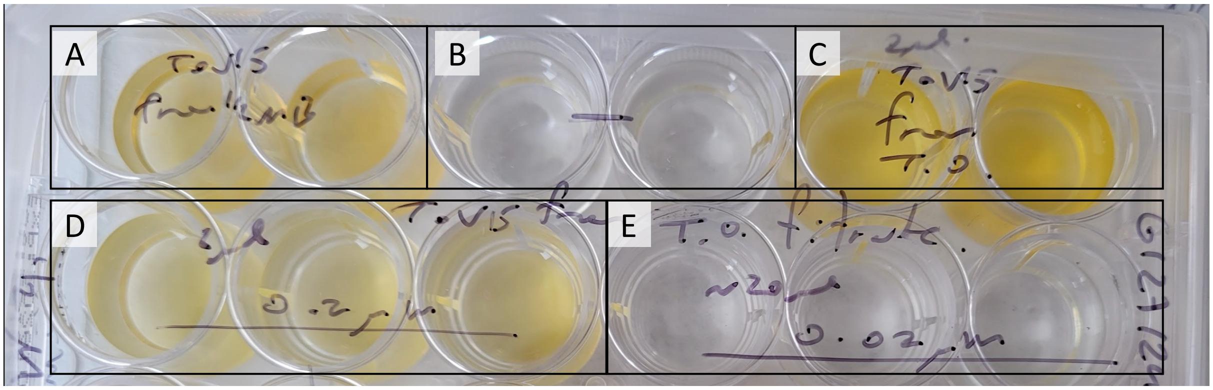


An example of a 24-well plate with 2 ml 50% MB in each well 2 weeks after *E. algicida* addition. **A.** *E. algicida* To15 from 50% MB. **B.** Control, addition of FSW. **C.** Addition of 2 µl co-culture of *T. oceanica* + *E. algicida*. **D.** Addition of 2 µl of < 0.2 µm fraction from *T. oceanica* + *E. algicida* co-culture. **E.** Addition of 20 µl of < 0.02 µm fraction from *T. oceanica* + *E. algicida* co-culture. Similar tests were conducted for all experiments with exudates.

### Figure S7 Effect of different media types on *T. oceanica* or *E. algicida* growth

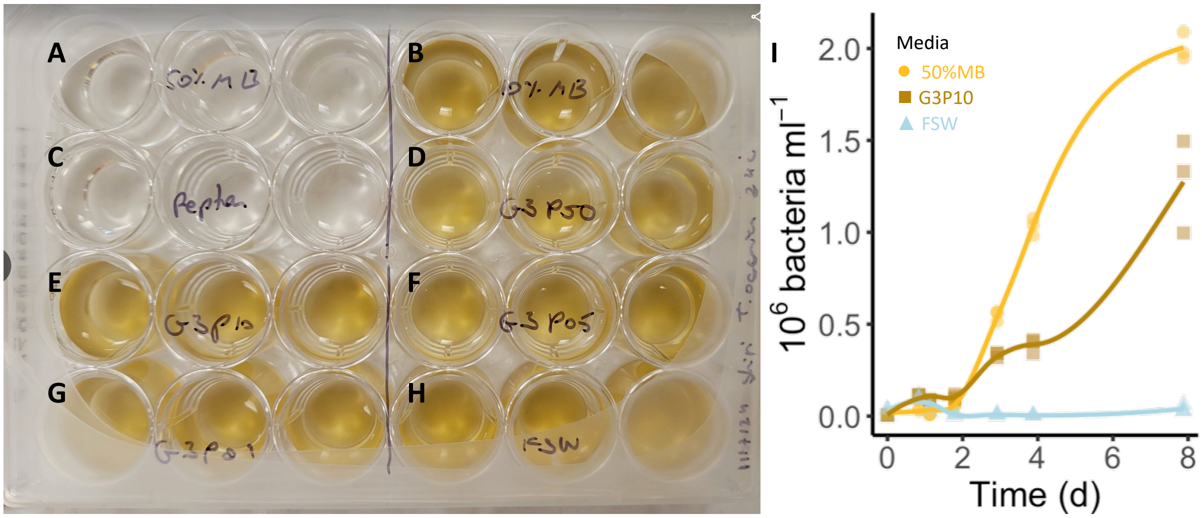


Image of *T. oceanica* cultures 7 days after addition of 10% (v/v) of various media. **A.** 50% MB. **B.** 10% MB. **C.** Peptone (300 mg ml$^{-1}$). **D-G.** Glucose (3 M) and varying concentration of peptone: **D.** peptone 150 mg ml$^{-1}$. **E.** peptone 30 mg ml$^{-1}$ . **F.** peptone 15 mg ml$^{-1}$ . **G.** peptone 3 mg ml$^{-1}$ . **H.** control- addition of FSW. **I.** *Ekhidna algicida* To15 growth as monoculture in different media types: 50%MB (yellow circles), 3M glucose + 30 mg ml$^{-1}$ peptone (G3P10, brown rectangles), filtered seawater without added nutrients (FSW, light blue triangles).
